# A genome-wide CRISPR screen identifies regulators of beta cell function involved in type 2 diabetes risk

**DOI:** 10.1101/2021.05.28.445984

**Authors:** Antje K Grotz, Elena Navarro-Guerrero, Romina J Bevacqua, Roberta Baronio, Soren K Thomsen, Sameena Nawaz, Varsha Rajesh, Agata Wesolowska-Andersen, Seung K Kim, Daniel Ebner, Anna L Gloyn

## Abstract

Identification of the genes and processes mediating genetic association signals for complex disease represents a major challenge. Since many of the genetic signals for type 2 diabetes exert their effects through pancreatic islet-cell dysfunction, we performed a genome-wide pooled CRISPR loss-of-function screen in human pancreatic beta cells. We focused on the regulation of insulin content as a disease-relevant readout of beta cell function. We identified 580 genes influencing this phenotype: integration with genetic and genomic data provided experimental support for 20 candidate type 2 diabetes effector transcripts including the autophagy receptor *CALCOCO2*. Our study highlights how cellular screens can augment existing multi-omic efforts to accelerate biological and translational inference at GWAS loci.

## Main

Genome-wide association studies (GWAS) have delivered thousands of robust associations for type 2 diabetes (T2D) and related glycemic traits but most map to non-coding regions and are presumed to have a regulatory function^1^. Incomplete fine-mapping efforts mean that most GWAS loci are not mapped to a single causal variant but rather include multiple variants in a credible set, each of which could potentially influence gene expression in a different cellular context. The typical step after fine-mapping involves attempts to connect the putatively causal variants – and the regulatory elements in which they reside – to the genes they regulate, using methods such as cis-eQTL colocalization, single cell chromatin co-accessibility, and DNA proximity assays (such as HiC)^2–5^. These approaches however have limitations. The two most important are the cell type- and context-dependency of these regulatory connections (which means that assays conducted in inappropriate cell-types or states may reveal variant-to-gene connections that have nothing to do with disease pathogenesis) and molecular pleiotropy (which means that variants of interest may regulate transcription of several genes in cis, obscuring the identity of the transcript responsible for disease mediation). These correlative approaches can generate hypotheses about candidate effectors but typically fall short of providing definitive evidence.

Perturbation studies can provide more compelling evidence of causation, but only if conducted in authentic models and using disease-relevant phenotypic readouts^6^. The strongest such evidence arises from disease-associated coding variants that provide a readout in free-living humans of the consequences of perturbations of gene and protein function, but the low frequency of most such variants limits this approach^2^. The availability of human cellular models and CRISPR based technologies provides an increasingly attractive alternative approach for generating genome-wide profiles of the phenotypic consequences of gene perturbation and thereby inform understanding of disease biology^6^. Central to this aspiration is confidence in the disease relevance of a cell type. For T2D, both physiological and epigenomic evidence highlight the central role of pancreatic islets, and consequently the insulin producing beta cells, in mediating disease risk ^7–11^. Substantial differences between mouse and human islets (architecture) and beta cells (ion channel composition, cell cycle regulation and glucose transporter expression) argue for the use of human tissue and cell lines^12–17^. We and others have generated large transcriptomic and epigenomic resources in this key human tissue enabling genome-wide integration of genetic and genomic data which has helped to identify candidate effector transcripts at T2D GWAS loci ^8, 11, 18, 19^. We now complement these resources with a genome-wide CRISPR loss-of-function (LoF) screen in the well characterized human pancreatic beta cell line EndoC-βH1 to identify genes which regulate insulin content ^20–22^. Using this resource, we provide cellular evidence to support the causal role of 20 genes at T2D loci in beta cell function and identify a novel role for the autophagy receptor *CALCOCO2* in human beta cell function.

## Results

### Development of a pooled CRISPR assay for human beta cell function

Glucose stimulated insulin secretion is the primary measure of beta cell function but is an unsuitable phenotypic readout for pooled high-throughput screening as secreted insulin cannot be linked directly to its cellular source. Changes in insulin content are also causally linked to T2D but unlike insulin secretion can be quantified using FACS.^23^ We therefore developed an assay for this closely-related and disease-relevant phenotype, endogenous intracellular insulin content, to enable fluorescence based cellular sorting using a primary monoclonal insulin antibody and a fluorescently labelled secondary antibody. Using this assay as our cellular phenotypic readout, we designed a pooled CRISPR LoF screening pipeline in the human pancreatic beta cell line EndoC-βH1 based on introducing single gene KOs through a lentiviral genome-wide CRISPR library followed by antibiotic selection to remove untransduced cells (Fig. 1a). Following gene knockout, cells were FAC-sorted into populations containing either low or high levels of intracellular insulin and the stably integrated sgRNAs within these cell populations identified through next-generation sequencing.

**Fig 1.**
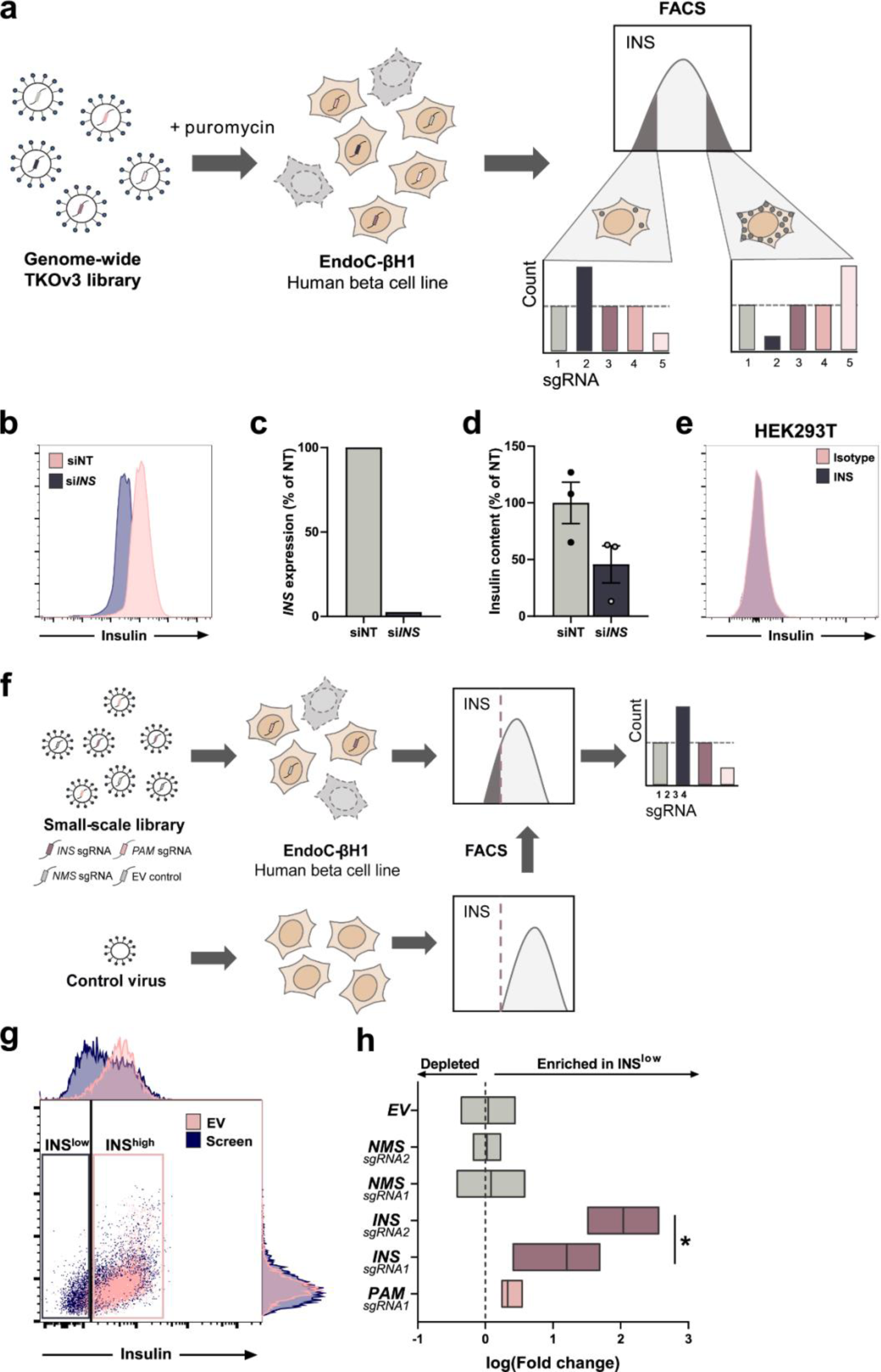
Optimization of a CRISPR screen for insulin content in EndoC-βH1. a) Pipeline for a genome-wide CRISPR loss-of-function screen in EndoC-βH1 from viral transduction (left) to antibiotic selection (middle) and a final FACS selection followed by sequencing and enrichment analysis of integrated sgRNAs (right). b) FACS staining for intracellular insulin in *INS* silenced EndoC-βH1 (si*INS*, blue) or their respective non-targeting controls (siNT, pink), c) the associated mRNA expression of *INS* and d) corresponding levels of intracellular insulin as measured by alphaLISA. e) FACS staining using the insulin antibody (blue) or isotype control (pink) in the human embryonic kidney cell line HEK293T. f) Pipeline for a small-scale CRISPR screen in EndoC-βH1 targeting only three genes *(NMS, INS* and *PAM)* alongside EV control cells. FACS sorting gates for low insulin (dashed line) were determined based on control cells transduced in parallel. sgRNA enrichment was assessed in INS^low^ compared to INS^high^. g) FACS staining in EndoC-βH1 comparing insulin staining of EV control cells (pink) and cells from the small-scale CRISPR screen (blue).Respective boxes indicate sorting gates INS^low^ and INS^high^. h) sgRNA enrichment for all individual sgRNAs in INS^low^ compared to INS^high^ from control (grey) samples and positive controls (*INS* and *PAM*, purple and pink). All data are presented as mean ± SEM from two (g,h) or three independent experiments and representative FACS plots are shown with their respective silencing efficiency. Data were analyzed using one (h) or two-sample t-test (d). NT, non-targeting. LFC, log2(fold change); EV, empty vector. P-value * < 0.05.

To ensure the FACS based insulin content readout in EndoC-βH1 was highly specific and sensitive, we chose an antibody and staining protocol based on comparing several complementary strategies (Supplementary Fig S1). Near complete siRNA-based knockdown of *INS* in EndoC-βH1 resulted in two populations with distinct average fluorescence intensities, validating the sensitivity of the INS antibody (Fig 1b,c). The level of separation was in concordance with an independent alphaLISA based quantification of insulin content demonstrating around 46% of residual protein after knockdown (Fig 1d). The specificity of the antibody was further confirmed in the non-*INS* expressing human embryonic kidney cell line HEK293T (Fig 1e). Finally, the screening protocol was validated by targeting three genes in a small-scale screen including the positive control genes *INS* and *PAM*, which are known to induce a reduction in insulin content upon deletion, and *NMS*, a negative control which is not expressed in EndoC-βH1^23^ (Fig 1f). Compared to an empty vector (EV) control coding for Cas9 only, the cells demonstrated, as expected, reduced INS FACS signal and enrichment of *INS* and *PAM* sgRNA within the INS^LOW^ population, whereas sgRNAs targeting *NMS* showed no difference (Fig 1g,h). Consistent with its lower effect on insulin content as shown in previous studies, sgRNAs targeting *PAM* demonstrated a highly replicable but lower enrichment compared to sgRNAs targeting *INS*, further confirming the suitability of the pipeline for genome-wide screening.

### A FACS-based CRISPR screen identifies regulators of insulin content

We performed two independent genome-wide screen replicates in the slow-growing (doubling in 7 days) human beta cell line, EndoC-βH1, using the Toronto KnockOut version 3.0 (TKOv3) CRISPR library targeting 18053 protein-coding human genes with four sgRNAs per gene^24^. Per replicate, a minimum of 700 million cells were lentivirally transduced at a low multiplicity of infection (MOI) with a library coverage of around 500 cells per guide, thus requiring more than 4 months to obtain sufficient cells for the screen. Cells were subjected to puromycin selection for seven days followed by fixation, permeabilization, staining and FACS sorting (Fig. 1a). Compared to EV (Cas9) control cells, the cell population transduced with the CRISPR library demonstrated a wider insulin signal distribution, particularly towards lower insulin, indicative of altered insulin content due to CRISPR KO mediated effects (Fig. 2a, Supplementary Fig S2). Two cell populations were collected, those with low insulin content (INS^LOW^) and those with high insulin content (INS^HIGH^), followed by DNA extraction, sgRNA amplification and sequencing (Supplementary Fig S3). We used the MAGeCK algorithm to compare sgRNA abundance and identify sgRNAs that were enriched or depleted in INS^LOW^ cells relative to INS^HIGH^ cells^25^. Enriched sgRNAs identify genes which are positive regulators leading to a reduction in insulin content upon gene KO, and depleted sgRNAs identify negative regulators associated with an increase in insulin content upon gene KO. To reduce the number of false positive hits, genes were only classified as screening hits if sgRNAs demonstrated consistent effects across replicates and met the stringent threshold (FDR<0.1) for at least two of the four sgRNAs in both independent screening replicates. A total of 580 genes fulfilled these criteria: we considered these as robust, reproducible screening hits and took them forward for further evaluation (Supplementary Table 1).

**Fig 2.**
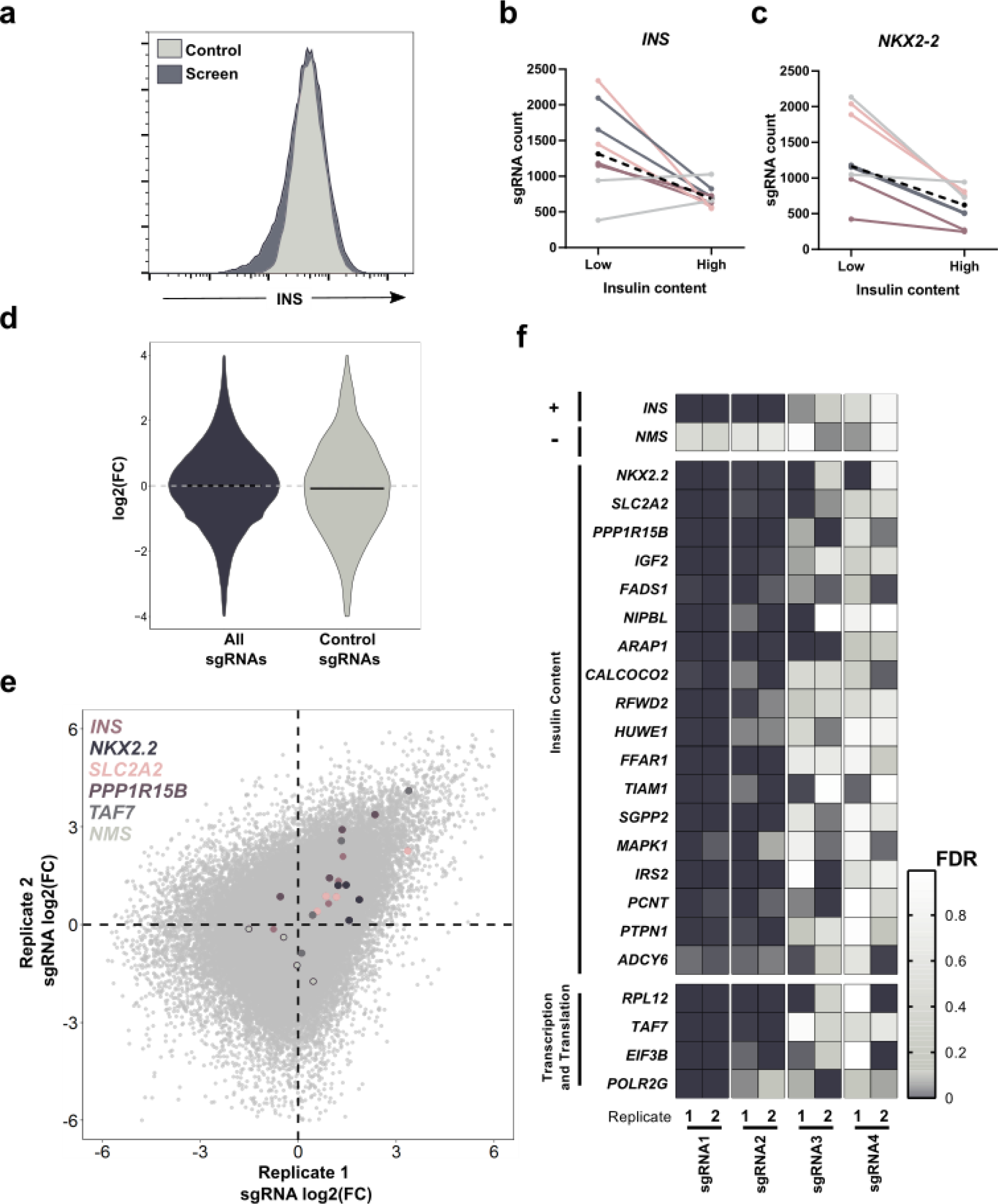
A CRISPR screen for insulin content in EndoC-βH1. a) Flow cytometry histograms showing insulin content in EndoC-βH1 transduced with the CRISPR screening library (blue) compared to cells transduced with EV virus (grey). b,c) Changes in sgRNA count from low to high insulin content screening sample with each colour representing the same sgRNA across the two screen replicates for (b) *INS* and (c) *NKX2-2*. The black dashed line represents the median sgRNA count for this gene. d) sgRNA distribution of log2(FC) for all sgRNAs (blue) compared to control sgRNAs targeting LacZ, EGFP and luciferase (grey). The black line indicates the median values for each group. e) sgRNA distribution of log2(FC) within each screening replicate with specific sgRNAs of interest (4 per gene) highlighted in individual colours per gene while overall sgRNA distribution is depicted in light grey. f) Individual genes of interest with their respective sgRNAs per replicate, the colour ranges from not significant (white) to highly significant (dark blue) based on the FDR and representing a significant sgRNA enrichment or depletion between INS^low^ and INS^high^. *INS* (+) and *NMS* (-) on the top panel represent positive and negative control genes, respectively. Data are from two independent genome-wide CRISPR screen replicates, screen FDR values were determined using MAGeCK (e,f) or through a two-sample t-test (d). Log2(FC), log2(fold change), EV, empty vector.

To evaluate the biological relevance of these screen hits, we asked if our screen was able to identify genes known to be involved in insulin regulation and beta cell function. As expected, sgRNAs targeting *INS* were enriched in the INS^LOW^ population, confirming the sensitivity of the screen (Fig. 2b,e,f). The overall enrichment of *INS* was lower than that of other known regulators of insulin secretion as only three out of the four sgRNAs induced an effect (Fig. 2b,f). Other established regulators of beta cell function and genes with known roles in monogenic forms of diabetes or T2D- risk were identified among the screening hits, such as the beta cell transcription factor *NKX2.2*, the glucose transporter *SLC2A2*, the cellular stress regulatory subunit *PPP1R15B* and the insulin-like growth factor *IGF2* (Fig. 2c,e,f) ^26–33^. Non-targeting control sgRNAs showed no effect on insulin content (Fig. 2d). In addition to direct regulators of insulin content, the screen also identified multiple general regulators of transcription and translation with indirect effects on the phenotype, such as the translation initiation factor *EIF3B*, the RNA polymerase II subunit *POLR2G*, the transcription coactivator *TAF7* and the ribosomal protein *RPL12* (Fig. 2f), all of them classified as common essential genes in non-beta cell CRISPR screens with longer culturing durations ^34–36^.

### Network analysis of CRISPR screening hits

We next considered whether the identified screening hits were enriched for functional classifications involved in the regulation of insulin content, by undertaking pathway and protein network analysis. Gene ontology (GO) term and Kyoto Encyclopedia of Genes and Genomes (KEGG) pathway analysis highlighted insulin and beta cell function associated categories such as ‘Type II diabetes mellitus’, ‘Insulin receptor signaling’ or ‘Calcium mediated signaling’, while transcription and translation related categories consistent with modifying total intracellular protein levels were also enriched (Fig. 3a)^37, 38^. STRING protein network analysis further emphasized the fundamental role of INS within the screen as a central node with several independent connections (Fig. 3b)^39^. The screening hits were significantly enriched for involvement in shared protein networks, providing additional confidence in the sensitivity of the screen to identify interrelated complexes (Fig. 3c, Supplementary Fig S4). Many of the hits mapped to the functional categories of ubiquitin mediated proteolysis and autophagy, mitochondrial ATP synthesis, vesicle trafficking and exocytosis, GPCR signaling, lipid metabolism and the MAPK signaling pathway, all of them containing both previously unknown regulators of insulin content and those with known roles in insulin secretion or beta cell function such as *RFWD2/COP1, HUWE1, FFAR1, TIAM1, SGPP2, MAPK1, IRS2, PCNT* and *PTPN1*^40–48^. The T2D associated gene *TBC1D4* mapped to a GTPase cluster and whilst the effects of this gene on T2D pathogenesis have been considered to be primarily associated with insulin resistance, the effect on insulin content observed in this screen is consistent with previous studies indicating a crucial role within beta cells^49–54^. The GTPase network also contained *ARAP1*, a gene previously linked to T2D susceptibility adding another dimension to the interesting story at this locus where deletion or overexpression in mice has failed to detect an impact on beta cell function pointing towards the nearby genes *STARD10* or *FCHSD2* as modulators of insulin secretion and mediators of disease risk at this GWAS locus^55–57^.

**Fig 3.**
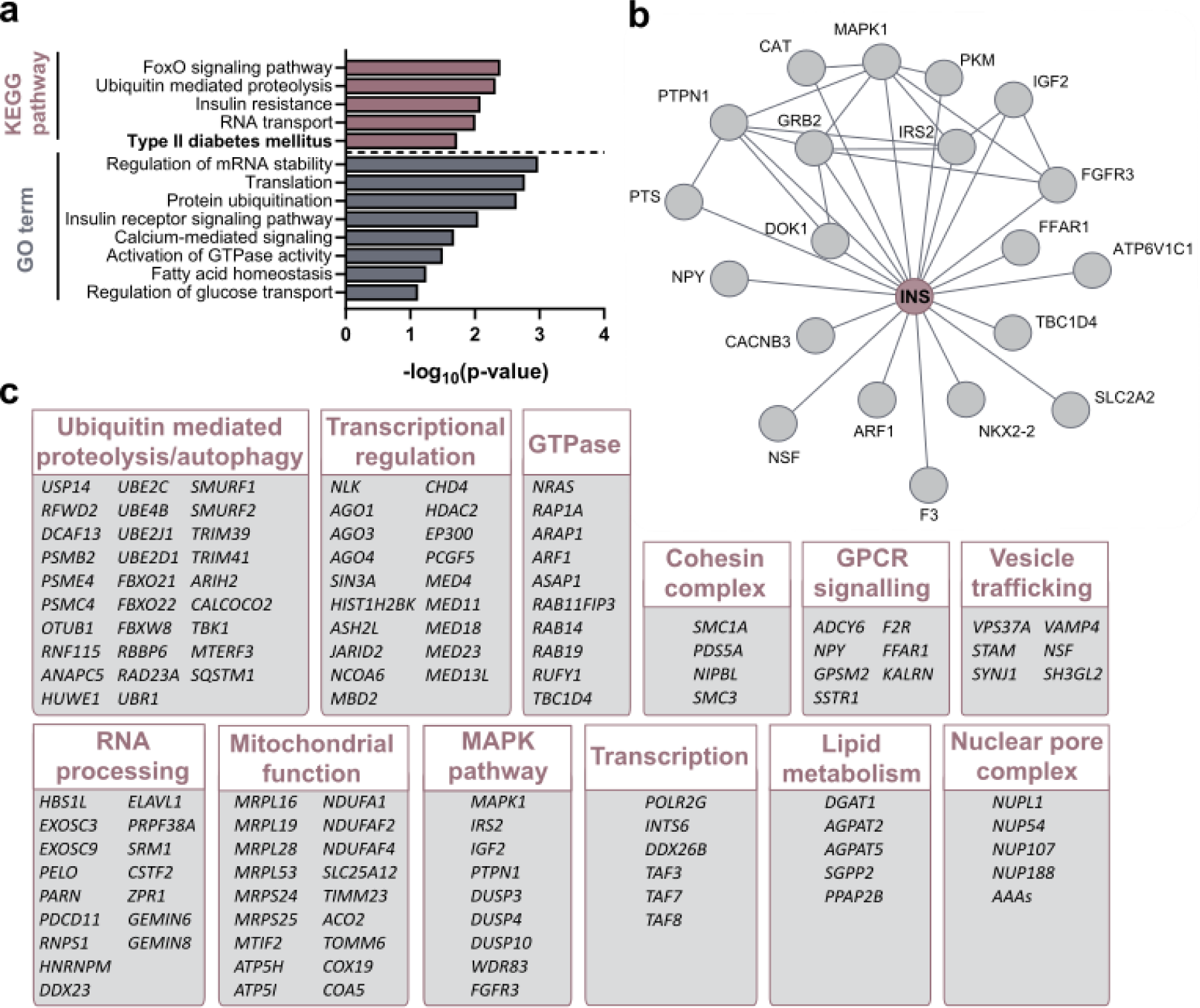
Integration of CRISPR screening hits. a) Pathway enrichment analysis assessing KEGG pathways or GO terms for biological processes. Selected pathways are shown, ranked by p-value. b,c) STRING pathway analysis showing protein-protein associations including physical and functional interactions b) between INS and other screening hits and c) for screening hits clustered into functionally associated groups.

### Integrative analysis of CRISPR screening hits with effector transcript prioritization methods

Next, we sought to use our cellular screen as complementary perturbation evidence to support the assignment of effector genes at T2D loci. We applied three complementary approaches which have been developed to combine genetic association results for T2D with diverse sources of genetic and genomic data and thereby generate lists of the “effector” gene(s) most likely to mediate the genetic association at each signal^58–60^. All methods are described and the predicted effector gene lists provided on the T2D Knowledge Portal (https://t2d.hugeamp.org) ^61^. The union of the effector gene assignments from the three methods generated a list of 336 candidate effector transcripts at T2D loci, with no candidates identified by all three methods. The intersection of this list with the set of screen hits, yielded 20 genes (3% of screening hits and 6% of predicted effector transcripts, Supplementary Table 2) for which our screen provided biological evidence for a role in a disease relevant phenotype. None of these 20 genes were identified by all 3 prediction methods. Of these 20 genes, 5 (25%) are assigned as “causal” by the curated *heuristic* effector gene prediction method with a further 4 (20%) assigned between strong, moderate and possible.

To compare how our perturbation screen performed against other commonly applied approaches for effector transcript prioritization, we explored, a relatively modest albeit the largest to date, eQTL study of 420 human islet donors^19^. In this study, colocalization between T2D-GWAS and islet cis-eQTL signals was supported by two methods (COLOC and RTC) at 20 loci, rising to 39 when statistical support for colocalization was required from only one of the approaches. Whilst both approaches identified similar numbers of genes, no gene was common between them, suggesting that the islet eQTLs are unlikely to manifest as reduced insulin content and additional cellular phenotypes (e.g. insulin secretion or ER stress) may underly their effect. These observations support the inclusion of multiple approaches for optimal effector gene prioritization and encourage both larger eQTL studies to detect further signals, which could then intersect with hits from this screen, and an expansion of targeted and/or genome-wide screens for additional cellular phenotypes (e.g. insulin secretion) across disease relevant cell types.

### Loss of the T2D associated gene *CALCOCO2* reduces insulin content

We focused our functional studies on the screening hit *CALCOCO2* (encoding Calcium Binding and Coiled-Coil Domain protein) on the basis that its role in the human beta cell has not been explored (Fig. 4a)^18, 62–65^. *CALCOCO2* is a ubiquitin-binding autophagy receptor and plays a crucial role in selective autophagy by linking the degradation target to the autophagic machinery^66, 67^. So far, the gene has been shown to initiate autophagy for invading pathogens (xenophagy) and damaged mitochondria (mitophagy)^66–68^. *CALCOCO2* maps to the T2D GWAS locus usually named for the gene *TTLL6,* but fine-mapping data highlighted a coding variant within *CALCOCO2* (rs10278, p.Pro347Ala) as the most likely causal variant^9^. Physiological clustering (based on patterns of association of the T2D-risk allele with other diabetes-related phenotypes) has revealed that this locus is likely to exert its primary effect on disease risk through the modulation of beta cell function^9^. In this CRISPR screen, KO of *CALCOCO2* resulted in consistent enrichment of *CALCOCO2* targeting sgRNAs in the low insulin content sample, corresponding to a decrease in insulin content (Fig. 4b).

**Fig 4.**
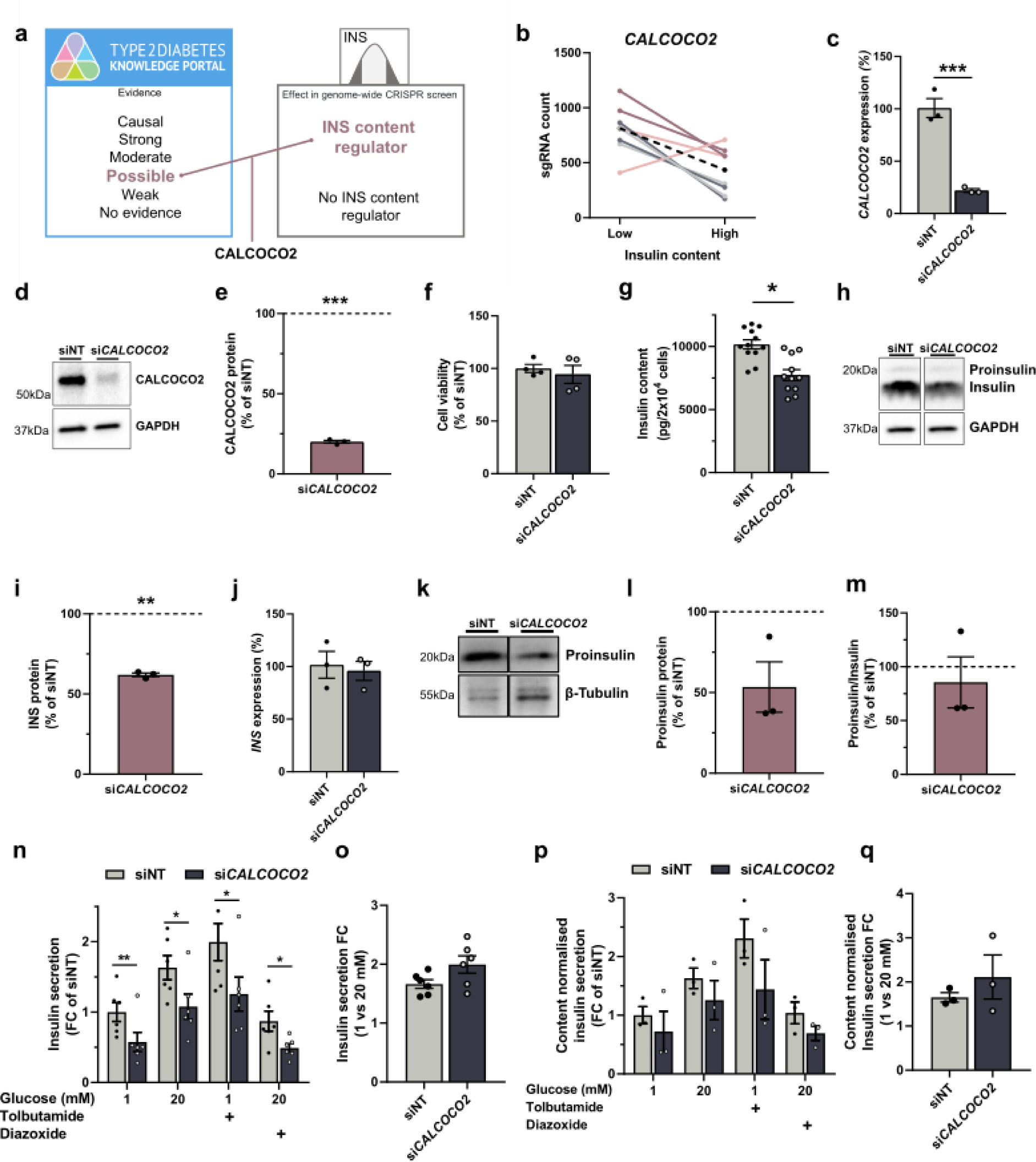
*CALCOCO2* knockdown reduces insulin content in EndoC-βH1. a) Gene prioritization approach combining genes with low evidence as T2D effector transcripts (as outlined in the integration approach) while being identified as a screening hit, highlighted *CALCOCO2.* b) Changes in sgRNA count from low to high insulin content sample with each color representing the same sgRNA across the two screen replicates. The black dashed line represents the median sgRNA count for this gene. c-m) All data are from si*CALCOCO2* treated EndoC-βH1 compared to non-targeting control cells. c) mRNA expression of *CALCOCO2*. d) Protein level of CALCOCO2 and its loading control GAPDH. e) Quantification of CALCOCO2 western blot data, normalized to GAPDH and siNT control cells. f) Cell count measurements. g) Insulin content in pg per 20 000 cells, measured by alphaLISA. h) Protein level of insulin and its loading control GAPDH. The antibody also detects proinsulin but only gives a weak signal compared to mature insulin. i) Quantification of insulin and western blot data normalised to GAPDH and siNT control cells. j) mRNA expression of *INS*. k) Protein level of proinsulin and its loading control β-Tubulin. l) Quantification of proinsulin western blot data normalised to β-Tubulin and siNT control cells. m) Ratio of proinsulin to insulin for western blot data. n,p) Insulin secretion normalised to siNT (n) or to insulin content and siNT (p) in 1 mM, 20 mM, 1 mM+100 μM tolbutamide or 20 mM glucose+100 μM diazoxide. o,q) Insulin secretion fold change from 1 to 20 mM glucose normalised to siNT (o) or to insulin content and siNT (q). The protein level is displayed as percentage of siNT which is highlighted as dotted line at 100%. All data are mean ± SEM from at least three independent experiments. Data were analyzed using two-sample t-test (c), one-sample t-test (e,i,l,m), one-way (f,g,j,o,q) or two-way (n,p) ANOVA with Sidak’s multiple comparison test. P-values * < 0.05, ** < 0.01, *** < 0.001. FC, fold change; NT, non- targeting.

None of the other genes at the *CALCOCO2* GWAS locus showed an effect on insulin content in the screen, further supporting *CALCOCO2* as the likely effector transcript at this T2D risk locus (Supplementary Fig S5).

We obtained independent confirmation of the effect of *CALCOCO2* loss on insulin content through siRNA-based silencing in EndoC-βH1. Silencing of *CALCOCO2* was highly efficient and induced a mean mRNA and protein reduction of 78.1% and 80.0%, respectively (Fig. 4c-e). Reduction of *CALCOCO2* did not affect cell viability (Fig. 4f). Intracellular insulin content was assessed using two independent antibody-based detection methods. Consistent with the results of the CRISPR screen and highlighting its sensitivity, insulin content was significantly reduced in *CALCOCO2* silenced cells by 24.0% and 38.0% as measured by an alphaLISA and western-blot based detection method, respectively (Fig. 4g-i). This reduction in insulin content was not due to decreased insulin gene expression nor was it due to altered proinsulin processing as the ratio of proinsulin to insulin was unaffected (Fig. 4j-m).

### *CALCOCO2* is required for human beta cell function

While we have shown that insulin content is reduced in EndoC-βH1 upon loss of *CALCOCO2*, altered glucose stimulated insulin secretion cannot be concluded based on changes in insulin content alone.

We therefore independently assessed the effects of *CALCOCO2* knockdown on insulin secretion in EndoC-βH1. Across different glucose and KATP modulator conditions (tolbutamide and diazoxide), insulin secretion was significantly reduced by 39.3% upon silencing of *CALCOCO2* (Fig. 4n). While total insulin secretion was affected, the glucose stimulation index from low to high glucose was unchanged, indicating an appropriate cellular response to changes in glucose concentrations (Fig. 4o). Normalization of insulin secretion to insulin content reduced the difference between *CALCOCO2* silenced cells and controls to 30.2%, indicating that the level of insulin content played a partial role in influencing the insulin secretory response (Fig 5p,q).

**Fig 5.**
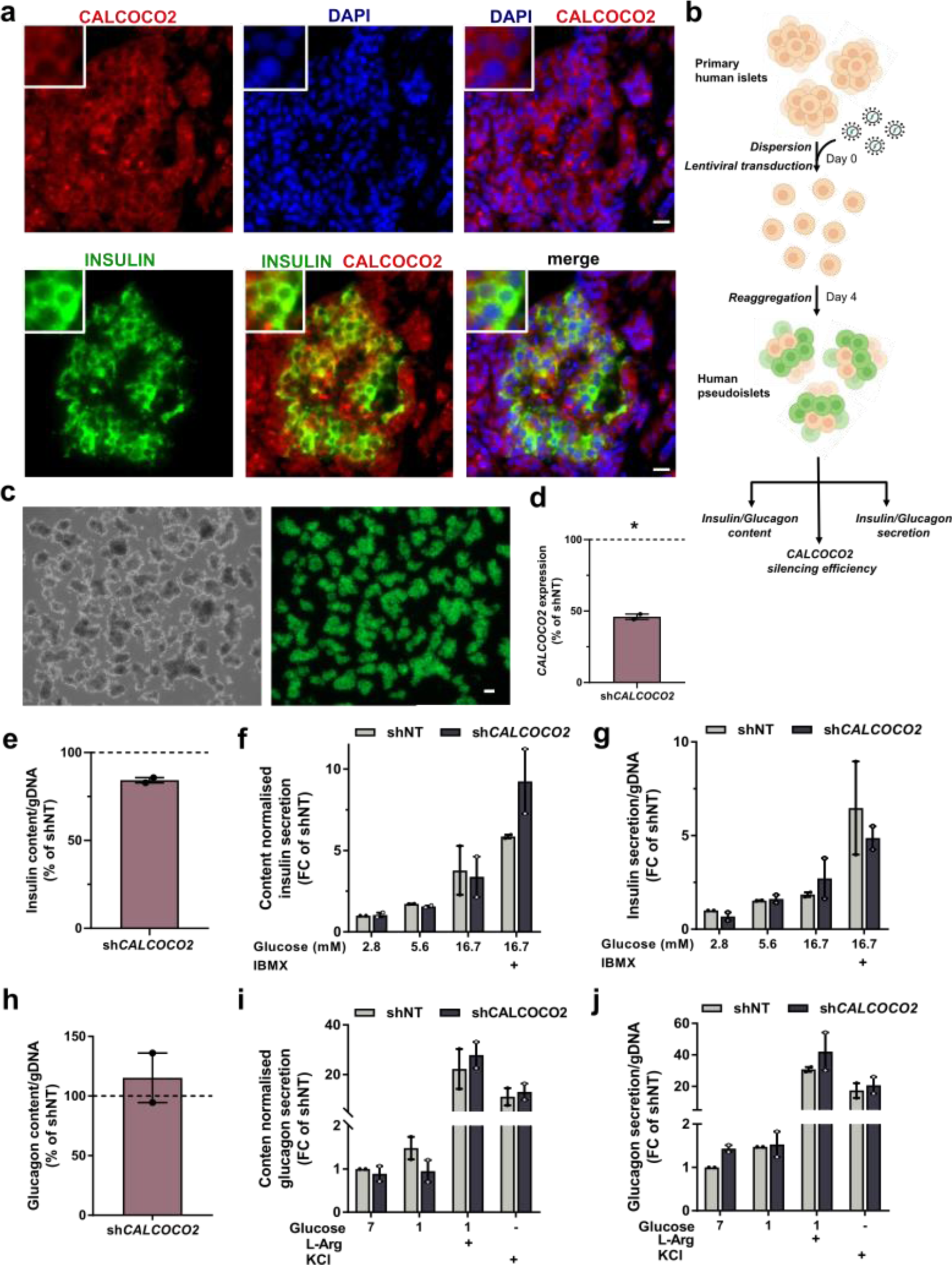
*CALCOCO2* knockdown reduces insulin content in primary human pseudoislets. a) Immunofluorescence staining of primary human islets in pancreas sections. Sections were double stained for INS (green) and CALCOCO2 (red). Cell nuclei were counterstained with DAPI (blue). Scale bar is 10 μm. b-j) All data are from sh*CALCOCO2* pseudoislets compared to non-targeting control pseudoislets (shNT). b) Pseudoislet transduction and formation from primary human islets. c) sh*CALCOCO2* pseudoislets under bright field (left) and GFP positive pseudoislets (right). Scale bar is 100 μm. d) mRNA expression of *CALCOCO2*. e,h) Intracellular insulin (e) and glucagon (h) content. f,g,i,j) Insulin (f,g) or glucagon (i,j) secretion normalised to content (f,i) or to genomic DNA (gDNA) (g,j) in 2.8 mM, 5.6 mM, 16.7 mM or 16.7 mM glucose + IBMX for insulin secretion and 7 mM, 1 mM, 1 mM glucose + L-Arginine or KCl for glucagon secretion. All data are mean ± SEM from two independent donor. Data were analysed using two-way ANOVA Sidak’s multiple comparison test (f,g,i,j) or one-sample t-test (d,e,h). P-values * < 0.05. FC, fold change; NT, non-targeting.

We extended our studies to determine *CALCOCO2* expression and localization in primary human pancreatic islets. The gene is expressed at a similar level as other critical beta cell genes such as *KCNJ11*, in both EndoC-βH1 and primary human islets^69^. Immunofluorescence studies in human pancreas sections confirmed protein localization to the cytoplasm of beta cells through colocalization with INS (Fig. 5a). Consistent with its ubiquitous role across cell types, exocrine ductal and acinar cells also stained positive for CALCOCO2 (Supplementary Fig S6).

To determine whether the effects of *CALCOCO2* loss can be replicated beyond *in vitro* models of human beta cells and in other secretory islet cell types, we performed shRNA knockdown of *CALCOCO2* in primary human islets from two independent human donors. Islets were dispersed, lentivirally-transduced with shRNA targeting *CALCOCO2* (sh*CALCOCO2*) and reaggregated to form pseudoislets (Fig. 5b). Successful transduction of pseudoislets was confirmed by assessing GFP co-expression (Fig. 5c). *CALCOCO2* expression was on average reduced by 54.0% compared to pseudoislets transduced with non-targeted control shRNA (shNT, Fig. 5d). In line with results in EndoC-βH, intracellular insulin demonstrated a modest but consistent reduction of 15.7% (Fig. 5e). If the lower silencing efficiency in pseudoislets is taken into account, insulin content is affected to the same extent in both the experimental model and primary human tissue. Upon incubation with a range of glucose concentrations, pseudoislets failed to demonstrate significantly affected insulin secretion (Fig. 5f,g). Stimulation with high glucose and the secretion potentiator IBMX increased secretion of insulin upon normalization to content in sh*CALCOCO2*, this effect was lost upon normalization to genomic DNA and therefore reflects the impact of changes in insulin content on secretion. We similarly assessed glucagon content and secretion to identify potential effects of *CALCOCO2* in alpha cells (Fig. 5h-j). While sh*CALCOCO2* showed some trend towards blunted glucagon response to glucose starvation, none of the secretory conditions or glucagon content were significantly changed upon *CALCOCO2* reduction. Together, our findings show that *CALCOCO2* regulates human islet beta cell function.

### The T2D-associated genes *QSER1* and *PLCB3* do not affect insulin content

To confirm that this screen correctly identified T2D associated genes which were likely to exert their effect on disease risk through modulating insulin content and beta cell function, we examined additional genes which have been predicted to be causal for T2D on the T2D Knowledge Portal but were not identified as screening hits (Fig. 6a)^59^. The prioritized genes *QSER1* and *PLCB3* both contain coding variants consistent with a causal role in T2D but their mechanism of effect or tissue of action has not yet been resolved ^9, 70^. *PLCB3* plays a role in the amplification pathway in the beta cell but has also been associated with insulin action in a multi-trait physiological clustering approach^9, 71, 72^. *QSER1* has recently been identified as a DNA methylation regulator associated with pancreatic differentiation defects^73^. In the screen, neither of these genes showed a consistent sgRNA depletion or enrichment in either insulin content populations and were not classified as hits (Fig. 6b,c). We again used siRNA knockdown in our targeted follow-up studies and reduced gene expression by 78.3% and 51.2% upon targeting of *QSER1* and *PLCB3*, respectively (Fig. 6d). Consistent with our negative screening results, knockdown of either gene did not affect intracellular insulin content (Fig. 6e). In addition, INS expression and insulin secretion under different glucose conditions or KATP modulator conditions were unchanged (Fig. 6f, Supplementary Fig S6). Together, this genome-wide CRISPR screen was able to correctly identify the insulin regulatory function of *CALCOCO2* whilst also distinguishing between cellular phenotypes related to islet-cell development (Q*SER1*) or insulin action (*PLCB3*).

**Fig 6.**
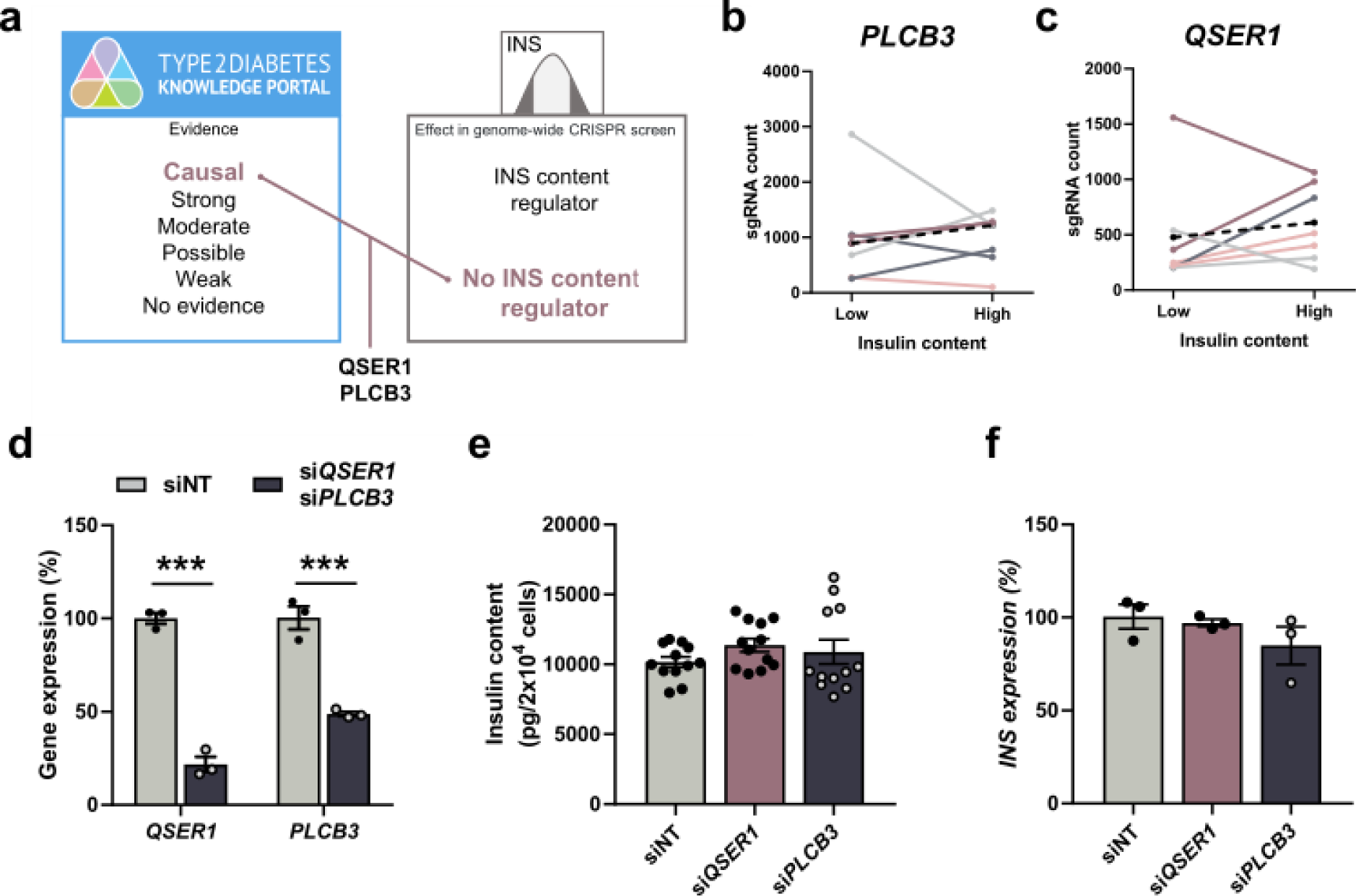
*QSER1* or *PLCB3* knockdown does not affect insulin content in EndoC-βH1. a) Gene prioritization approach combining genes with causal evidence as T2D effector transcripts (as outlined in the integration approach) while also not being a screening hit, highlighted *QSER1* and *PLCB3*. b) Changes in sgRNA count from low to high insulin content sample with each color representing the same sgRNA across the two screen replicates. The black dashed line represents the median sgRNA count for this gene. d-f) All data are from si*QSER1* or si*PLCB3* treated EndoC-βH1 compared to non- targeting control cells. d) mRNA expression of *QSER1* and PLCB3 in their respective silenced cells (blue) compared to siNT control cells (grey). e) Insulin content in pg per 20 000 cells. f) mRNA expression of *INS*. All data are mean ± SEM from three independent experiments or two independent genome-wide CRISPR screen replicates. Data were analysed using two-sample t-test (d) and one-way ANOVA with Sidak’s multiple comparison test (e, f). P-values *** < 0.001. FC, fold change; LFC, log2 (fold change); NT, non-targeting.

## Discussion

Recent efforts to identify effector transcripts at GWAS association signals have concentrated on the integration of disease relevant transcriptomic and epigenomic datasets or focused on detailed mechanistic studies at a single locus ^8, 11, 18, 19, 23, 63, 74–77, 78^. In most cases these studies are unable to distinguish between pleiotropic effects of variants on regulatory elements and gene expression which are inconsequential for the disease phenotype in question. It is increasingly recognized that genome-wide perturbation datasets in disease relevant cell types could close the gap between effector transcript and disease relevant biology and thus enable focused translational efforts. For T2D, there is substantial evidence to support a central role for defects in pancreatic islet-cell dysfunction along with an understanding of disease relevant cellular phenotypes thus opening the possibility to generate relevant genome-wide cellular perturbation datasets. Here, we present a genome-wide pooled CRISPR loss-of-function screen applying a functionally relevant FACS readout to perform a comprehensive characterization of regulators of insulin content in the human beta cell line EndoC-βH1.

Our pooled genome-wide CRISPR screening approach was successful in identifying robust modulators of insulin content as shown through the detection of not only the insulin gene (*INS)* itself but also genes involved in monogenic types of diabetes or known regulators of insulin transcription and secretion such as the transcription factor *NKX2.2* or the glucose transporter *SLC2A2*. Through the integration of our screening hits with the results from three different currently available effector gene prioritization tools, 20 genes were identified as effector transcripts working through beta cell dysfunction. Our data add to a growing number of genome-wide multi-omic datasets which can be harnessed to close the gap between genetic discovery and biological mechanisms and serve as a model for future cellular studies not only in human beta cells but also in other disease relevant cell types.

Furthermore, our unbiased genome-wide data will be relevant for other traits/diseases where the pancreatic beta cell plays a role, including type 1 diabetes.

We initially identified the T2D risk associated gene *CALCOCO2* as a CRISPR screening hit and a novel positive regulator of insulin content. We selected this gene for further study on account of its potential to underlie a T2D GWAS signal at the *CALCOCO2* locus and due to a lack of knowledge on its role in beta cell biology. Our focused functional studies using complementary siRNA-based knockdown not only confirmed its effect on insulin content but also revealed a role in insulin secretion. We therefore provide, for the first time, functional data connecting *CALCOCO2* to beta cell function, a likely effect on disease pathogenesis and tissue of action, expanding the evidence for a causal role beyond its genetic coding association with T2D risk ^9, 70, 79^. Detailed mechanistic studies will be the focus of future investigations but *CALCOCO2*’s role as a selective autophagy receptor hint at defective insulin secretion because of dysregulated autophagy. *CALCOCO2* is an essential receptor to induce Parkin-mediated mitophagy (the degradation of dysfunctional mitochondria) and the observed effect on insulin secretion is consistent with other studies such as for the mitophagy regulator *Clec16a* ^68, 80–83^. In addition, the mitophagy regulators *SQSTM1* and *TBK1* were also identified as screening hits, further highlighting the relevance of this cellular pathway. In addition to its association with T2D risk based on GWAS and its intracellular role, *CALCOCO2* has also been implicated in studies of the plasma proteome, demonstrating that reduced circulating levels of *CALCOCO2* are associated with increased T2D risk ^84, 85^. Plasma proteins are key druggable targets and disease biomarkers, positioning *CALCOCO2* at the center of investigations focusing not only on understanding the underlying functional mechanism of T2D but also potential therapeutic or diagnostic strategies ^86^.

Our CRISPR screen in the human beta cell line EndoC-βH1 identified more than half of all protein networks that were also identified in a previous CRISPR screen assessing insulin content in a mouse insulinoma cell line, highlighting conserved interspecies regulatory networks ^87^. While shared protein networks were identified in both CRISPR screens, only a small proportion of screening hits (7% or 43 genes) were overlapping, highlighting the importance of performing the screen in a human cellular model system but also potentially diverging experimental strategies. Of the 20 genes from our CRISPR screen that were overlapping predicted effector transcripts, only one gene and the primary phenotypic readout in both screens, *INS*, was also identified by the CRISPR screen in the mouse insulinoma cell line ^87^.

The reason many investigators select to work in rodent beta cells lines is one largely of experimental ease. Although the human beta cell line EndoC-βH1 demonstrates functional characteristics closely resembling human primary cell it is also associated with difficult growth and culture characteristics ^20–22^. To obtain perturbation evidence in this physiologically relevant cellular model, we overcame the associated obstacles and variations across passages through highly consistent culturing conditions, integration of two independent replicates, stringent analysis parameters focusing on high reproducibility and rigorous proof of concept studies.

This is the first step towards the generation of comprehensive genome-wide perturbation assays in disease relevant human cell lines for T2D and represents a proof of concept that cellular screens can augment genomic efforts linking variants to regulatory elements and transcripts to bridge the gap between gene expression and disease relevant cellular biology. This genome-wide effort in a human cell line should accelerate efforts for further screens, both loss and gain of function, for additional disease relevant cellular phenotypes in authentic human models.

In summary, we have developed and performed a genome-wide pooled CRISPR screen in a model of human beta cells, providing a comprehensive perturbation database to associate genes of interest with a direction of effect, tissue of action and functional mechanism. We have successfully applied the screening hits as a prioritization tool for causal genes at T2D GWAS loci and highlight *CALCOCO2* as a novel modulator of insulin secretion and beta cell function with a likely causal role in T2D.

## Methods

### Cell culture

HEK293T were routinely passaged in DMEM 6429 (Sigma-Aldrich) containing 10% fetal calf serum, 100 U/ml penicillin and 100 µg/ml streptomycin. EndoC-βH1 were routinely passaged in 5.5 mM glucose containing growth medium (Sigma-Aldrich) as previously described^88^. All cells were mycoplasma free (Lonza) and grown at 37°C and 5% CO2.

### Human islet procurement

Deidentified human islets and pancreas samples were obtained from three healthy, nondiabetic organ donors procured through the Integrated Islet Distribution Network (IIDP) and the Alberta Diabetes Institute Islet Core (donor information in Supplementary Table 3).

### Cloning of individual sgRNAs

Single CRISPR sgRNAs were cloned into plentiCRISPRv2 (Addgene #52961) as previously described^20, 89^. Briefly, BsmBI compatible tails (5’CACCGX3’ and 5’AAACYC3’) were added to complementary sgRNA oligonucleotides and annealed (Supplementary Table 4). PlentiCRISPRv2 was digested using BsmBI (Fermentas), ligated with annealed sgRNA oligonucleotides using Quick Ligase (NEB) followed by transformation and amplification in Stbl3 competent cells (Invitrogen). Correct sgRNA integration was validated through Sanger Sequencing and plasmids were further used to produce lentivirus.

### Pooled sgRNA library amplification

Toronto human knockout pooled library (TKOv3) containing 71,090 based on a lentiCRISPRv2 backbone was a gift from Jason Moffat (Addgene #90294). The library was transformed and amplified using 25 μl Endura Competent Cells (Lucigen) and 100 ng plasmid libraries per electroporation reaction as previously described^24^. Transformation efficiency was determined through serial dilution to ensure sgRNA representation and plasmids were subsequently extracted using Plasmid Mega kits (Qiagen). Library representation was confirmed through sequencing on a NextSeq500 (Illumina) using 75 base pair (bp) single end reads.

### Lentiviral production and transduction

Lentivirus for individually cloned sgRNAs and pooled libraries was produced and titered as previously described^20^. Briefly, CRISPR plasmids were co-transfected in 60 % confluent HEK293T with packaging vectors pMD2.G (Addgene #12259), psPAX2 (Addgene #12260) using jetPRIME transfection reagents (Polyplus transfection). Viral supernatant was collected at 48 h and 72 h post transfection and concentrated using ultracentrifugation. The functional viral titer was determined in EndoC-βH1 through measuring the percentage of survival after transduction with different viral dilutions and antibiotic selection. Cell viability was assessed using the CyQUANT Direct Cell Proliferation assay (Invitrogen). The required amount of lentivirus to achieve 26 % survival which is representative of an MOI of 0.3 was calculated and used in subsequent small-scale or genome-wide screen transductions. EndoC-βH1 were transduced for 6 h and selected for 7 days in 4 μg/ml puromycin with media changes as required.

### Genome-wide CRISPR screen in EndoC-βH1

Two independent genome-wide CRISPR screens were performed consecutively in EndoC-βH1 with independent lentiviral CRISPR library transductions, FACS sorting and sgRNA sequencing. Each screen was performed at a coverage of 500 cells per sgRNA and a MOI of 0.3. A total of 670 million and 744 million cells were transduced in replicate one and two, respectively. The cells were transduced as described, incubated in 4 μg/ml puromycin for 7 days with media changes as required to remove dead cells followed by collecting, staining and FACS analysis.

### INS intracellular staining and fluorescence-activated cell sorting

EndoC-βH1 cells were harvested and incubated with LIVE/DEAD Fixable Far Red Dead Cell Stain (Thermo Fisher) for 30 min at room temperature to distinguish live from dead cells and washed in 1% BSA in PBS. The cells were fixed and permeabilized using the BD Cytofix/Cytoperm kit (BD Biosciences) for 20 min at 4°C and washed using Perm/Wash Buffer (BD Biosciences). Staining with primary antibodies was performed overnight at 4°C followed by incubation with suitable secondary antibodies diluted in Perm/Wash Buffer (Supplementary Table 4). The samples were filtered through a 70 μm cell strainer and sorted on a FACSAria III (BD Biosciences) using a 100 μm nozzle. Isotype and transduction controls stained with each antibody alone were analyzed alongside the samples. INS^LOW^ and INS^HIGH^ samples were gated based on live and single cells with lower INS levels compared to EV control cells (INS^LOW^) and the highest 10 % of INS-expressing cells (INS^HIGH^). Sorted cell samples were stored at -80 °C until DNA extraction. Flow cytometry data was analyzed using Flowjo 10.6 (BD Biosciences).

### Preparation of genomic DNA for next-generation sequencing

DNA was extracted from frozen FACS sorted samples using QIAamp Blood Maxi/Mini Kit (Qiagen) and processed for sequencing in a two-step PCR approach. Integrated sgRNAs were amplified using Q5 polymerase (NEB) with 2.5 μg DNA input per reaction and an optimized number of cycles per sample according to manufacturer’s instructions. Specific Illumina TruSeq adapters were attached to each sample using Q5 polymerase (NEB) with an optimized number of cycles (Supplementary Table 4). PCR products were run on a 2 % gel and purified using QIAquick Gel extraction kit (Qiagen).

Each Illumina library sample was qPCR-based quantified using the KAPA Library Quantification Kit (Roche) prior to pooling and multiplexed sequencing on a NEXTSeq500 (Illumina) as 75 bp reads with standard Illumina sequencing primer and PhiX to approximately 20% spike-in.

### Analysis of pooled CRISPR screen

Raw fastq sequencing read files were merged for each sample using the “cat” command. To identify enriched and depleted sgRNAs in this CRISPR screen, we used the previously described MAGeCK (v0.5.9.2) algorithm^25^. Briefly, the “count” command was used to extract and count sgRNA sequences from the raw fastq files based on the TKOv3 library input yielding a mean read count of 981 aligned reads per sgRNA (Supplementary Fig S3). sgRNAs with low read counts of less than 10 and sgRNAs mapping to genes that were not expressed in EndoC-βH1 were excluded. The “test” command was used with the paired module to assess and analyse sgRNA enrichment or depletion between INS^LOW^ and INS^HIGH^ populations. The analysis median normalizes read counts across samples to account for varying read count depth and applies a mean-variance model to identify significant sgRNA enrichment or depletion. MAGeCK multiple-testing adjusted sgRNA-level enrichment scores were the basis for gene-level hit selection. We applied additional stringent criteria to only prioritise hits with highly consistent effects across both replicates requiring genes to have an FDR<0.1 for at least two out of their four sgRNAs in both independent replicates. CRISPR screen analysis was performed in Python 3.8 and R 3.5.

### Gene set enrichment and pathway analysis

Enriched GO and KEGG pathways within the screening hits were determined using the Database for Annotation, Visualization and Integrated Discovery (DAVID) with homo sapiens as list and background input^37^. Protein connectivity networks based on physical and functional interactions were identified among screening hits using STRING v11^39^. Only interactions with a high confidence score of ≥0.9 were selected.

### Gene silencing in EndoC-βH1

Forward siRNA-based silencing in EndoC-βH1 was performed at 24 h after plating using ON-TARGET plus SMARTpool siRNAs for *CALCOCO2, QSER1, PLCB3* or a non-targeting control (Dharmacon) at 15 nM final concentration. siRNAs were preincubated with 0.4% Lipofectamine RNAiMAX (Invitrogen) in Opti-MEM reduced serum-free medium (Gibco) for 15 min at room temperature to form transfection complexes before dropwise addition to the culture. Cells were assayed or harvested 72 h post transfection.

### CALCOCO2 silencing in primary human islets

Lentiviral constructs coding for shRNAs targeting human *CALCOCO2* were obtained from Dharmacon and virus was produced as described before. Primary islets were dispersed into single cells by enzymatic digestion (Accumax, Invitrogen) and transduced with 1 × 10^9^ viral units/1 mL lentivirus. Transduced islet cells were cultured in ultra-low attachment well plates for five days prior to further analysis.

### Gene expression analysis

RNA for gene expression analysis from EndoC-βH1 was extracted using TRIzol reagent (Invitrogen) and synthesized into complementary DNA using the GoScript Reverse Transcriptase System (Promega), both according to manufacturer’s instructions. RNA for gene expression analysis from primary human pseudoislets was extracted using the PicoPure RNA isolation kit (Life Technologies) and cDNA was synthesized using the Maxima first strand cDNA synthesis kit (Thermo Scientific), according to manufacturer’s instructions. Quantitative qPCR (qPCR) was performed using TaqMan real-time PCR assays on a 7900HT Fast Real-Time PCR System (all Applied Biosystems). Ct-values were analysed using the ΔΔCt method and target genes were normalized to the combined average of the housekeeping genes peptidylprolyl isomerase A (PPIA), glycerinaldehyd-3-phosphat-dehydrogenase (GAPDH) and TATA-box binding protein (TBP). *CALCOCO2* expression in EndoC- βH1 and primary islets was extracted from previously published and analyzed RNA sequencing data^69, 90^.

### SYBR Green based quantitative PCR

sgRNA integrations in the small-scale screen were quantified using SYBR Green based quantitative PCR and primer targeting the respective sgRNA sequences (Supplementary Table 4). DNA from FACS sorted EndoC-βH1 was extracted as described. Each sample was prepared using 20 ng of total DNA and SYBR Green PCR Master mix (Bio-Rad), following manufacturer’s instructions. The samples were amplified and analysed as described for gene expression experiments.

### Insulin secretion assay in EndoC-βH1

Silenced EndoC-βH1 were starved overnight at 48 h post transfection in culture medium containing 2.8 mM glucose followed by 30 min incubation in 0 mM glucose the next morning. Insulin secretion was initiated through static incubations in the indicated glucose or secretagogues conditions for 1 h. Insulin containing supernatant was harvested and adherent cells were lysed in ice-cold acid ethanol to release intracellular insulin. Insulin was quantified using the Insulin (human) AlphaLISA Detection Kit and the EnSpire Alpha Plate Reader (both Perkin Elmer) based on 1:10 and 1:200 dilutions for supernatant and insulin content, respectively. Secreted insulin was normalized to the level of intracellular insulin content or cell count, which was measured prior to cell lysis using the CyQUANT Direct Cell Proliferation assay (Invitrogen).

In vitro insulin and glucagon secretion assays in primary pseudoislets Batches of 25 pseudoislets were used per donor for *in vitro* secretion assays as previously described ^91^. For insulin secretion assays, pseudoislets were incubated at 2.8, 5.6, 16.7 and 16.7 mM + IBMX glucose concentrations for 60 min each and supernatants were collected. For glucagon secretion assays, pseudoislets were incubated at 7, 1, 1 mM +10 mM Arginine glucose concentrations for 60 min each and supernatants were collected. Human insulin in the supernatants and pseudoislet lysates were quantified using a human insulin ELISA kit (Mercodia). Human glucagon in the supernatants and pseudoislet lysates were quantified using a human glucagon ELISA kit (Mercodia). Secreted insulin and glucagon levels are presented as a percentage of gDNA, quantified from the pseudoislet lysates.

### Western Blot analysis

Whole-cell protein extracts were obtained from cell pellets through lysis in RIPA buffer (50 mM Tris pH 7.4, 150 mM NaCl, 1% Triton X-100, 0.5% sodium deoxycholate, 0.1% SDS) containing 1x protease inhibitor cocktail (Roche). Protein concentrations were quantified using the DC protein assay (Bio-Rad), 10 μg of total protein were mixed with sample buffer (4x Laemmli Buffer (Bio-Rad), 10% β-mercaptoethanol (Sigma-Aldrich)) and boiled for 10 min at 80°C. Denatured samples were run on a 4-20% Criterion TGX Stain-Free Precast Gel for 15 min at 300 V in Tris–glycine–SDS buffer and transferred to a 0.2 μm polyvinylidene difluoride (PVDF) using the Trans-Blot Turbo Transfer System (all Bio-Rad). Unspecific antibody binding was blocked through incubation with 3% BSA in Tris-buffered saline containing 0.1% Tween-20 (TBST) for 1 h at room temperature. Primary antibody incubations were performed at 4°C overnight followed by incubation with HRP-conjugated secondary antibodies for 1 h at room temperature (Supplementary Table 4). Blots were imaged using the ChemiDoc MP Imaging System (Bio-Rad) and reprobed as described using loading control antibodies of appropriate size. Image Lab 6.0 software (Bio-Rad) was used to quantify protein bands and proteins of interest were normalized to their respective loading control on the same blot.

### Immunofluorescence analysis

Human pancreatic tissue was obtained from subjects in post-mortem examinations in line with local and national ethics permission. Pancreata were fixed in 4% paraformaldehyde overnight, cryoembedded and sections of 4 μm were prepared. Tissue sections were dipped in distilled water, boiled for 20 min in target retrieval solution (Dako), permeabilized for 10 min at room temperature using 1% PBS-Triton X-100 (Sigma-Aldrich) and unspecific antibody binding was blocked using 1% BSA (Roche), 0.2% non-fat milk, 0.5% Triton-X 100, 1% DMSO in PBS (all Sigma-Aldrich).

Primary antibody incubations were performed overnight at 4°C followed by incubation with secondary Alexa Fluor-conjugated antibodies at room temperature for 1 h (Supplementary Table 4). The slides were mounted using VectaShield mounting media containing DAPI (Vector Laboratories) and imaged on a Zeiss AxioM1 (Zeiss) microscope using an x20 and x40 objective.

### Statistics

Statistical analyses were performed in Prism 8.1 (GraphPad Software). The number of biological independent replicates are shown as individual data points and the error bars represent the standard error of the mean. If appropriate, fold changes were plotted but statistical analysis was performed on log-transformed values. Normally distributed variables were compared between two or more groups using two-sample Student’s t-test or one-way analysis of variance (ANOVA) followed by Sidak’s multiple comparison test, respectively. Samples that were normalized to their respective control within a single replicate were analyzed using a one-sample Student’s t-test.

### Data availability

Fastq sequencing files from the CRISPR screen have been deposited in the European Nucleotide Archive (ENA) at EMBL-EBI. The accession number will be made available upon publication.

## Acknowledgments

A.L.G. is a Wellcome Senior Fellow in Basic Biomedical Science. A.K.G and S.K.T. are Radcliffe Department of Medicine Scholar. A.L.G. is funded by the Wellcome (200837) and National Institute of Diabetes and Digestive and Kidney Diseases (NIDDK) (U01-DK105535; U01-DK085545, UM1DK126185), the Stanford Diabetes Research Center (NIDDK award P30DK116074). We thank past and current members of the Gloyn lab for advice and encouragement, Dylan Jones for library amplification, Ruddy Montandon for FACS sorting and the Oxford Genomics Centre (Wellcome Centre for Human Genetics) for sequencing. We thank Yan Hang, Mollie Friedlander and the Islet Core of the Stanford Diabetes Research Centre for support with hormone secretion assays. We gratefully acknowledge organ donors and their families, and islet procurement through the Alberta Diabetes Institute Islet Core, Integrated Islet Distribution Program (U.S. NIH UC4 DK098085), the National Disease Research Interchange, and the International Institute for the Advancement of Medicine.

## Author contributions

A.K.G., S.K.T. and A.L.G. conceived the study. A.K.G., E.N-G. and S.K.T. performed CRISPR screening optimization experiments. A.K.G., E.N-G., R.B. and S.N. performed the CRISPR screen. E.N-G. and R.B performed sequencing library preparation. A.K.G. analyzed the CRISPR screening data. A.W-A contributed to the CRISPR screening analysis. A.K.G. and A.L.G. performed data integration analysis. A.K.G. performed the characterization of *CALCOCO2, QSER1* and *PLCB3* in beta cells. A.K.G. and R.J.B performed islet immunostainings. R.J.B., A.K.G. and V.R. performed islet shRNA knockdown experiments. S.K.K, D.E. and A.L.G. supervised the research. A.K.G. and A.L.G. wrote the manuscript. All authors approved the final draft of the manuscript.

## Supplementary Information

**Supplementary Fig S1.**
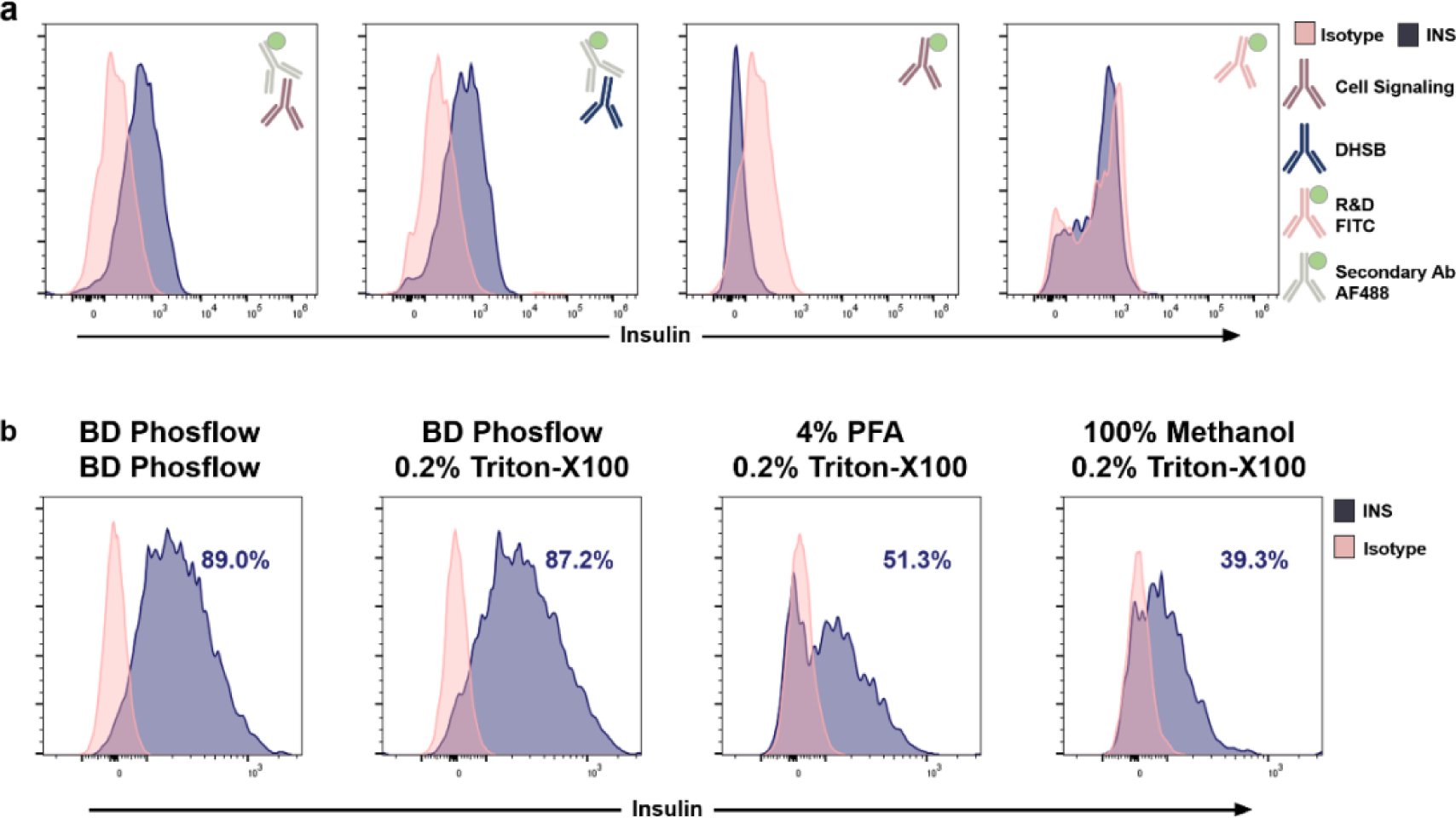
FACS optimization for intracellular INS staining. Histograms showing different FACS staining strategies in EndoC-βH1 comparing intracellular insulin (blue graph) and respective isotype controls (pink graph) using a) different monoclonal primary antibodies as indicated (purple, blue, pink antibody) in combination with a secondary Ab (grey) conjugated to AF488 (green) or direct conjugation to FITC (green) and b) using different staining protocols with various fixation (top row) or permeabilization buffers (bottom row) as indicated. Three independent experiments were performed and representative histograms are shown.

**Supplementary Fig S2.**
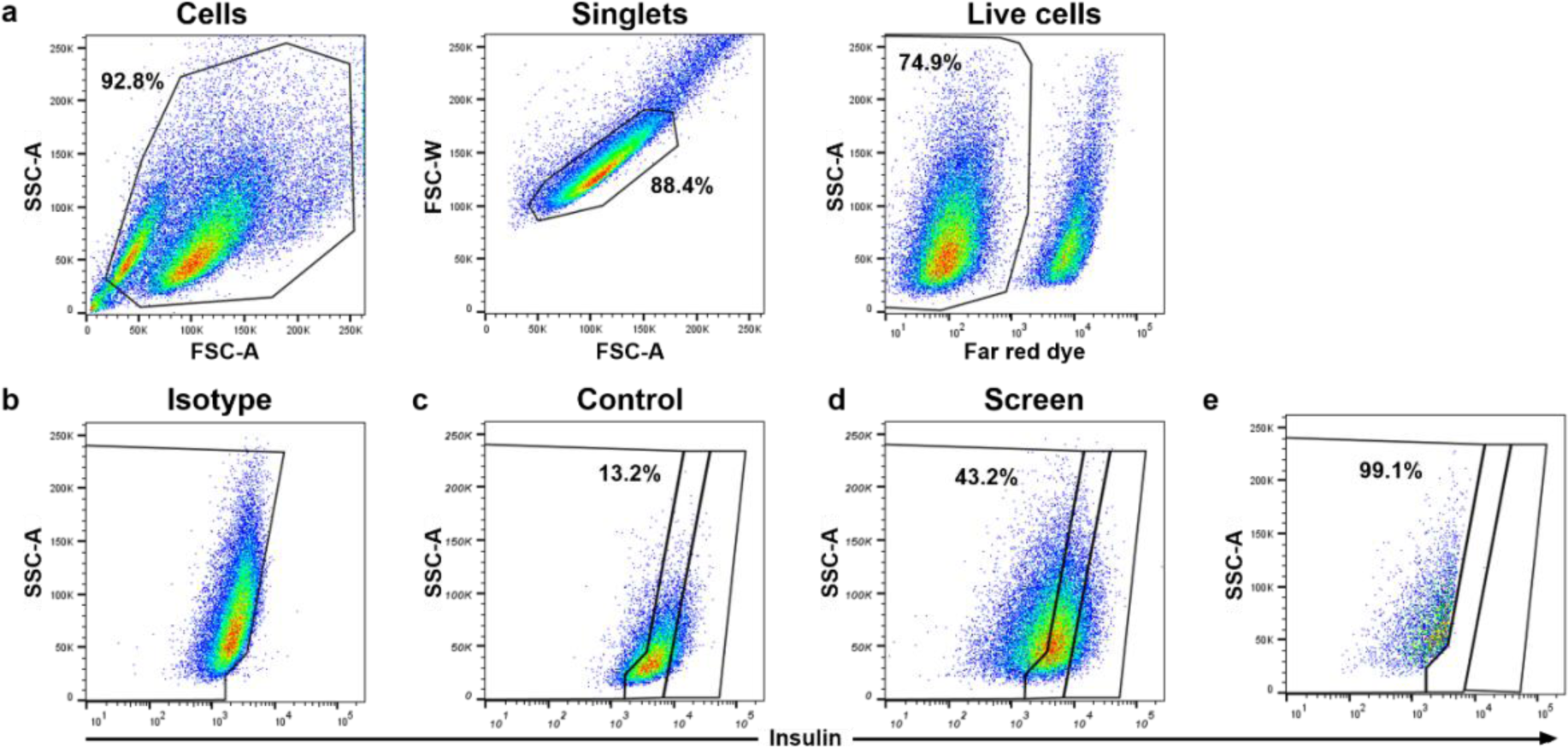
FACS sorting of the genome-wide CRISPR screen. Representative FACS images and sorting gates for stained EndoC-βH1 in the genome-wide CRISPR screens. a) Sorting gates assessing cell size, granularity and viability to exclude debris, define singlets and live cells. b-d) Insulin staining of the isotype control (b), control cells (c) and screening cells (d). e) Resorting of low insulin cell population to assess the sample purity. SSC-A, side scatter area; FSC-A, forward scatter area; FSC-W, forward scatter width.

**Supplementary Fig S3.**
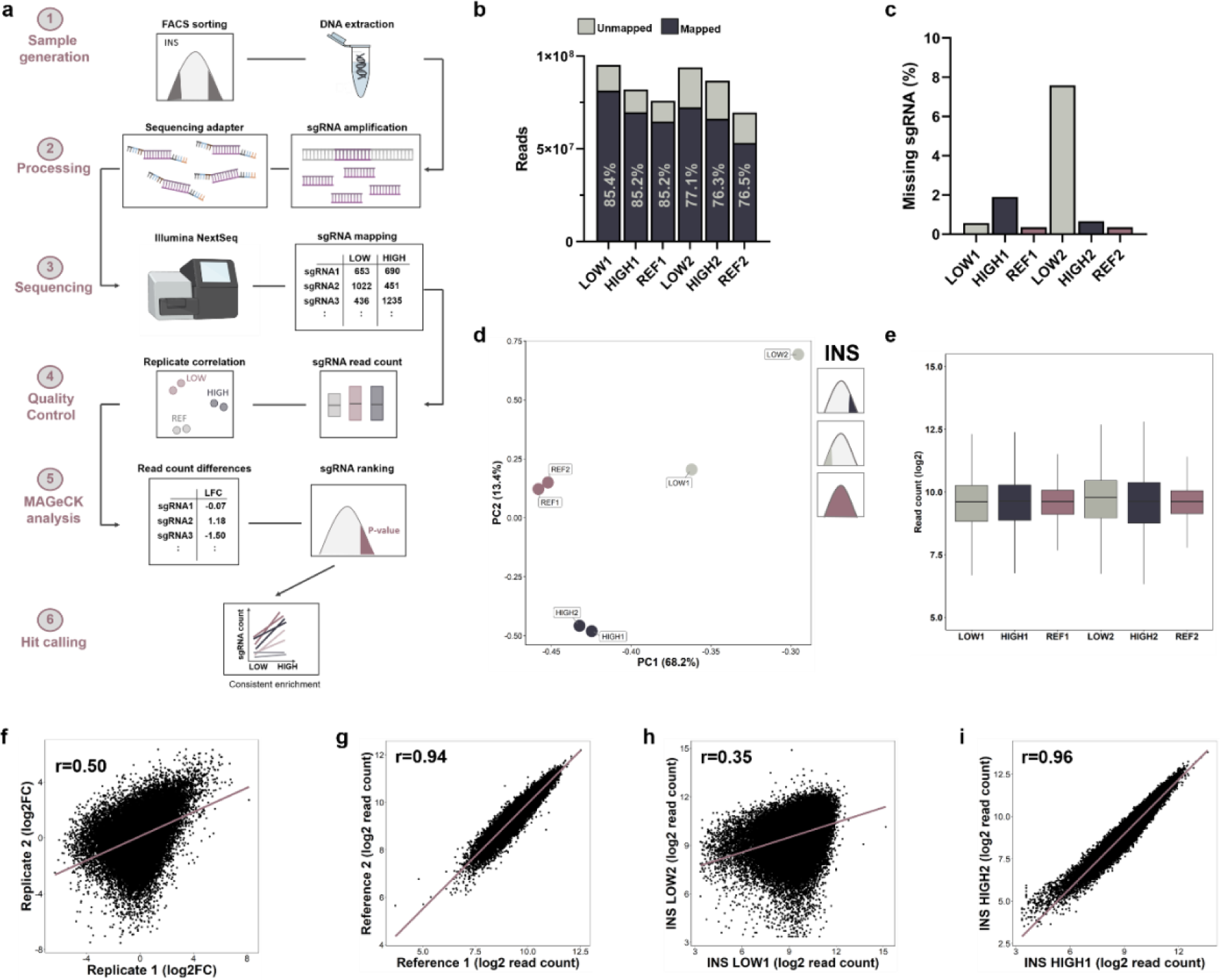
CRISPR screen outline and quality control. a) CRISPR screen outline from initial FACS sorting of transduced EndoC-βH1 to final sequencing analysis. b-c,e) Read count and mapped sgRNA quality control for low insulin (LOW1, LOW2), high insulin (HIGH1, HIGH2) and reference samples (REF1, REF2) from both replicates including: (b) total read counts and the proportion of mapped reads and (c) the number of zero-count sgRNAs as proportion of total sgRNAs and (e) normalized read count distributions. d) Principal Component Analysis for the first two components of low *(grey)* and high *(blue)* insulin content and reference samples *(purple)* from both replicates. f-i) Pairwise Pearson correlation analysis of fold changes between low and high insulin content for each replicate (f), reference replicates (g), low insulin content replicates (h) and high insulin content replicates (i). The purple line indicates the linear regression line. Log2FC, log2 (fold change); r, Pearson correlation coefficient.

**Supplementary Fig S4.**
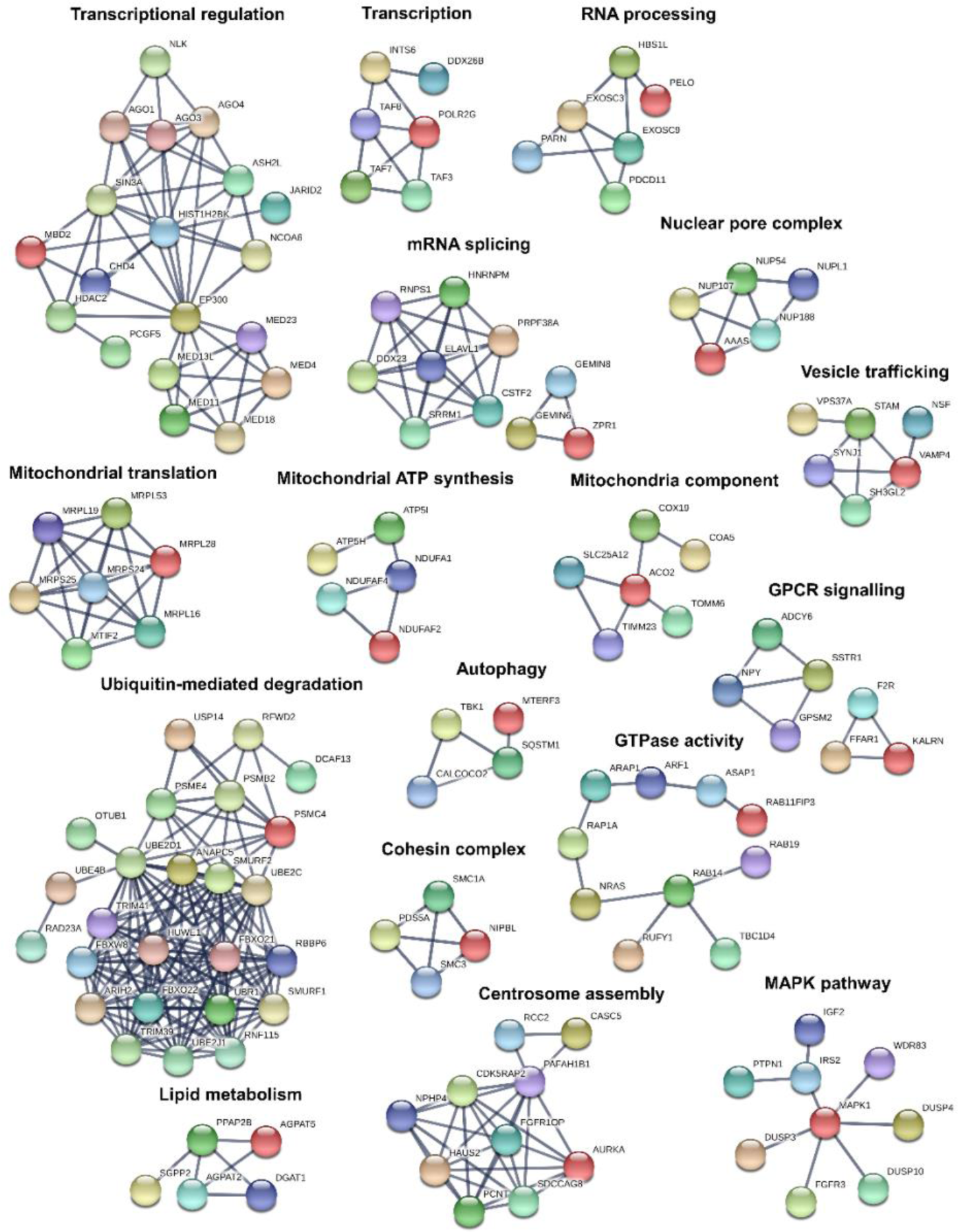
Protein association networks of screening hits. STRING pathway analysis showing protein-protein associations including physical and functional interactions for CRISPR screening hits. Clusters were manually identified, annotated, and individually plotted and selected clusters are shown. Confidence level to determine interactions was set to highest confidence (0.9).

**Supplementary Fig S5.**
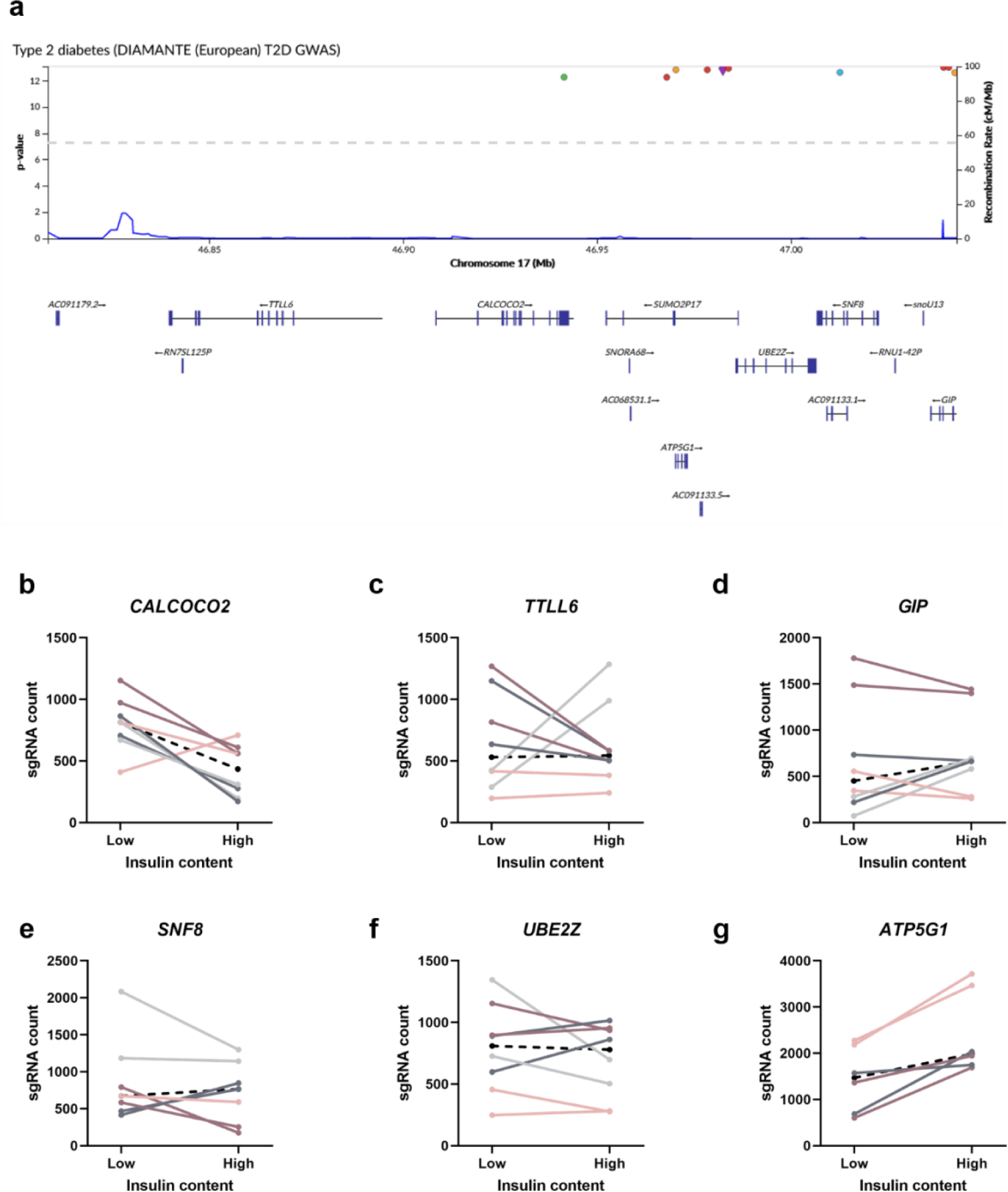
*CALCOCO2* locus with its candidate genes. a) *CALCOCO2* locus with credible set GWAS association signals *(top)* and neighbouring genes *(bottom).* b-g) sgRNA count distribution for all 8 sgRNAs per gene (4 sgRNAs per replicate) in the genome-wide CRISPR screen for genes at the *CALCOCO2* locus. Changes in sgRNA count from low to high insulin content sample with each colour representing the same sgRNA across the two screen replicates. The black dashed line represents the median sgRNA count for this gene.

**Supplementary Fig S6.**
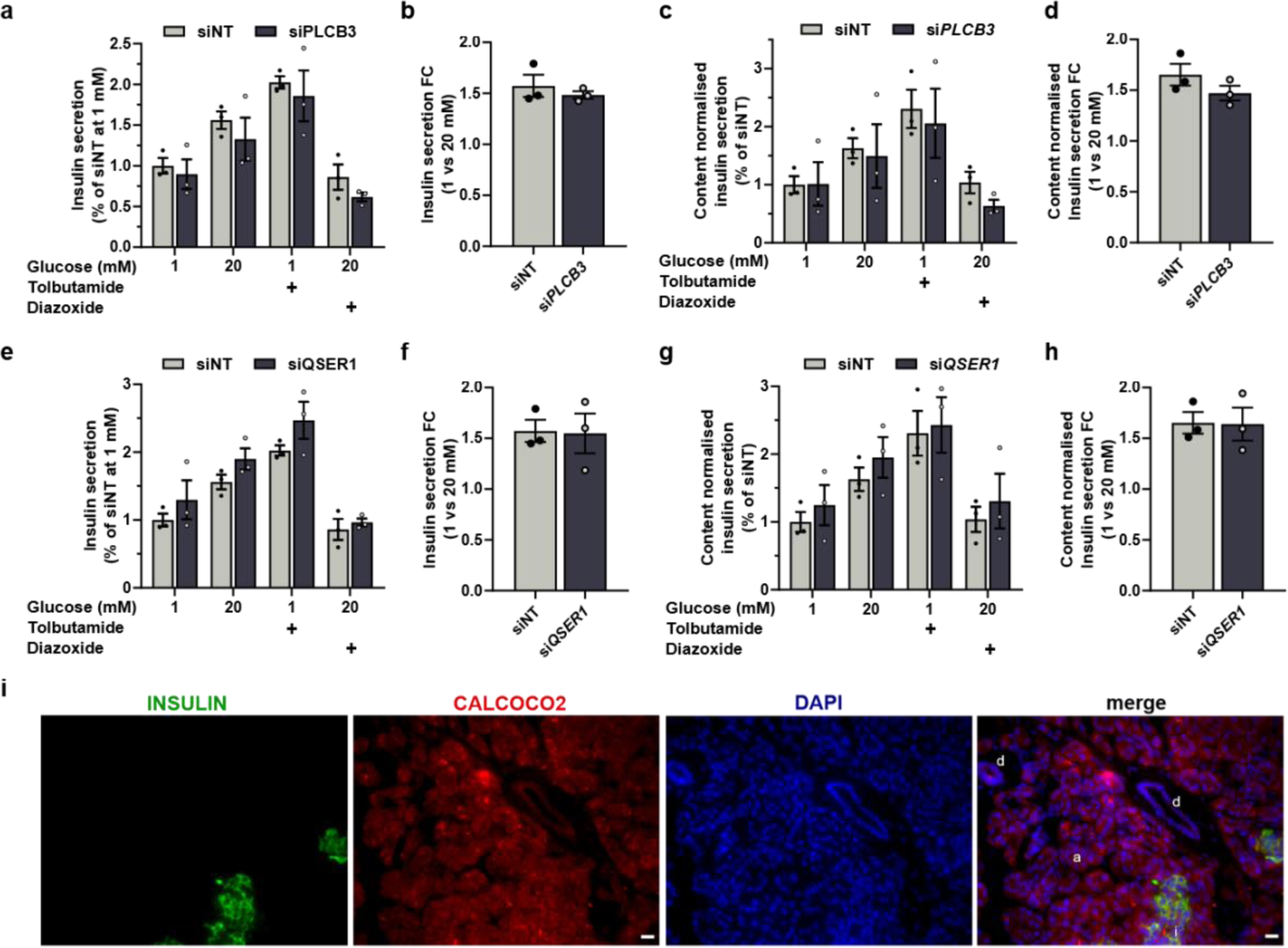
*QSER1* and *PLCB3* in EndoC-βH1, CALCOCO2 in the exocrine pancreas. a-h) All data are from siRNA treated EndoC-βH1 (blue) as indicated compared to non-targeting control cells (grey). (a,e) Insulin secretion normalised to siNT or (c,g) to insulin content and siNT in 1 mM, 20 mM, 1 mM+100 μM tolbutamide or 20 mM glucose+100 μM diazoxide. (b,d,f,h) Insulin secretion fold change from 1 to 20 mM glucose. All data are mean ± SEM from three independent experiments. Data were analysed using two-way ANOVA Sidak’s multiple comparison test (a,c,e,g) and two-sample t-test (b,d,f,h). P-values *** < 0.001. FC, fold change; NT, non-targeting. i) Immunofluorescence staining of exocrine tissue in pancreas sections. Sections were double immunostained for INS (*green*) and CALCOCO2 (*red*). Cell nuclei were counterstained with DAPI (*blue*). Scale bar is 10 μm. d: ductal cells; a: acinar cells; i: islet.

Supplementary Table 1 – CRISPR Screening Hits.

A complete list of the 580 genes meeting the stringent criteria set for reproducibly effecting insulin content levels in the genome-wide CRISPR screen. Genes are shown based on fold change and the direction of effect (increase or decrease) on insulin content indicated.

**Supplementary Table 2.** Prioritized causal genes for T2D based on integration with T2D effector gene predictions.

| **Genome-wide CRISPR screen** | **eQTL for T2D (Viñuela et al. 2020)** |
| --- | --- |
| *AGPAT2* | *ADCY5* |
| *ARAP1* | *AP3S2* |
| *CALCOCO2* | *CAMK1D* |
| *DCUN1D4* | *STARD10* |
| *FADS1* | *CEP68* |
| *IGF2* | *DGKB* |
| *INS* | *GPSM1* |
| *IRS2* | *HMG20A* |
| *MCCC1* | *PLEKHA1* |
| *MED23* | *ITGB6* |
| *NKX2-2* | *RNF6* |
| *NUDT3* | *TCF7L2* |
| *PELO* | *UBE2E2* |
| *PPP1R15B* | *NKX6-3* |
| *RBMS1* | *GRB14* |
| *SHQ1* | *HAUS6* |
| *SIN3A* | *IGF2BP2* |
| *SLC2A2* | *KLHL42* |
| *TBC1D4* | *ABCB9* |
| *ZNF101* | *SCD5* |
|  | *SLC12A8* |
|  | *SLC7A7* |
|  | *PDE8B* |

Common genes from effector transcript predictions and the genome-wide CRISPR screen (left) or from a recent human islet eQTL studies for T2D (right)<sup>19</sup>.

**Supplementary Table 3.** Human Tissue Donor Details.

| **Tissue source** | **Sample Identifier** | **Sex** | **Age (yrs)** | **BMI (Kg/m<sup>2</sup>)** |
| --- | --- | --- | --- | --- |
| IIDP | SAMN18021384 | Male | 56 | 24.2 |
| IIDP | SAMN18479112 | Male | 44 | 24.4 |
| ADI | R398 | Female | 51 | 25.4 |
| Other | OD30535 | Female | 31 | 48.2 |

IIDP, Integrated Islet Distribution Program (https://iidp.coh.org). ADI, Alberta Diabetes Institute http://www.bcell.org/adi-isletcore.html

**Supplementary Table 4.**
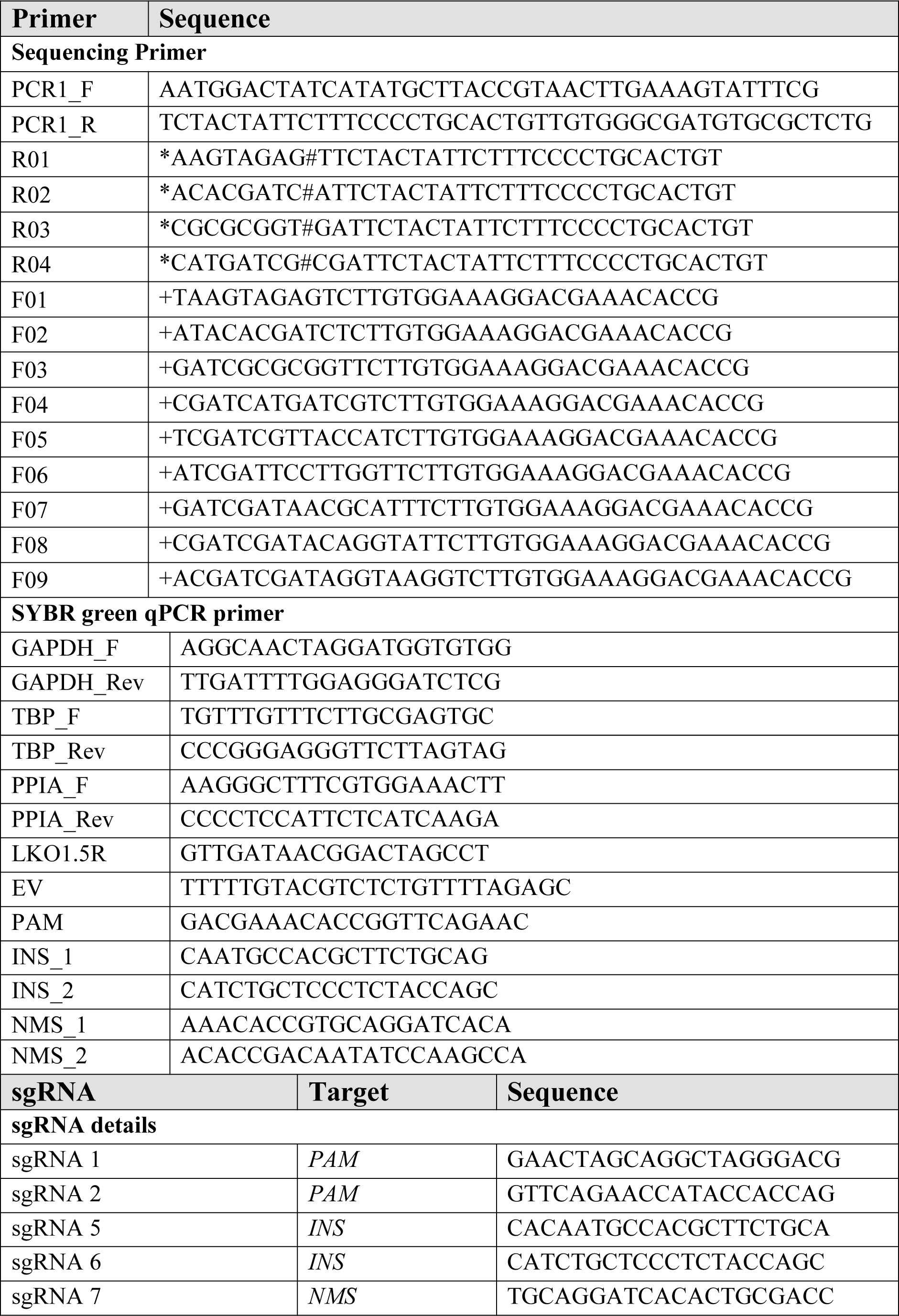
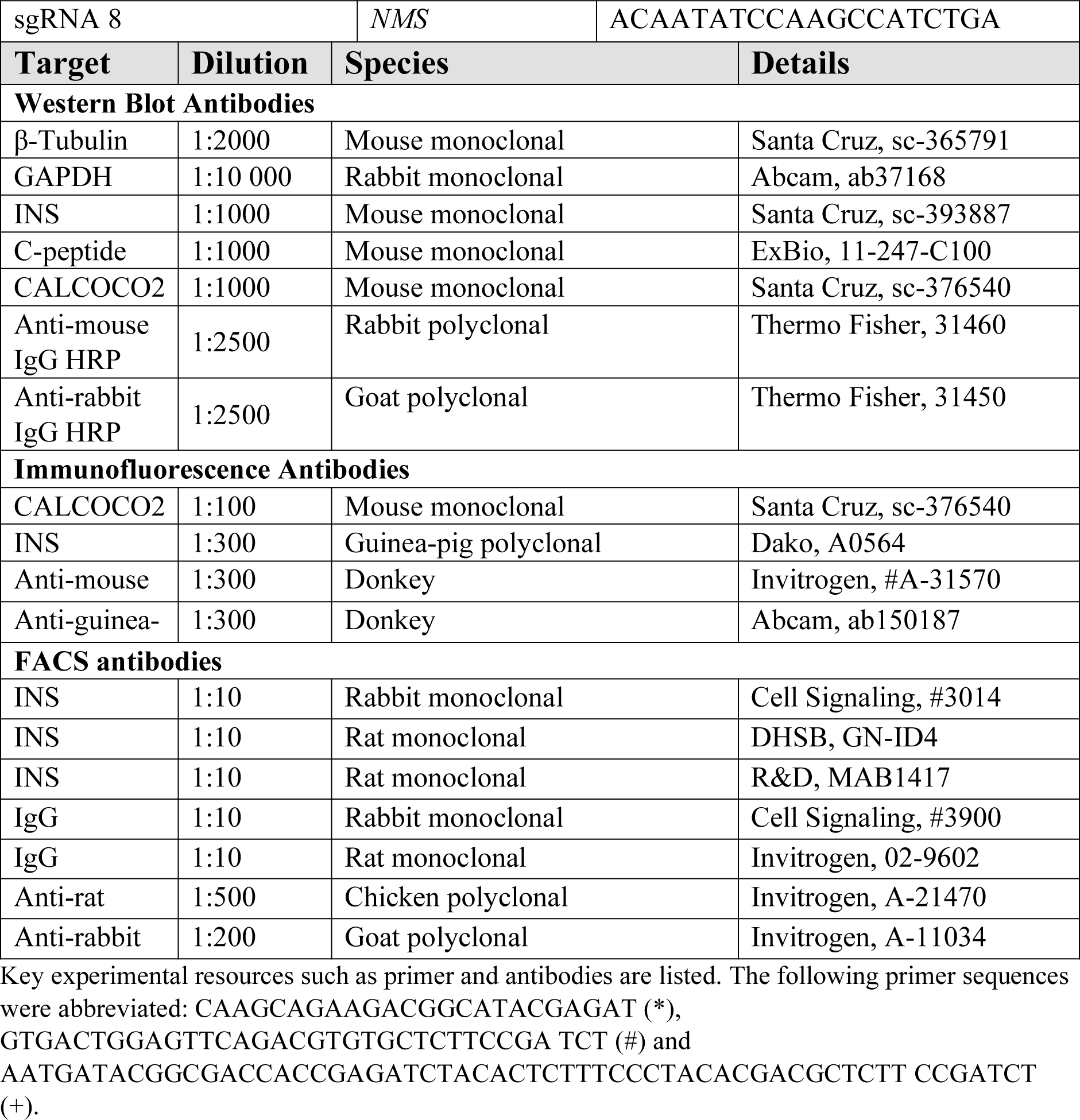
Resource Table.

