## Supplementary Table 1 for "A genome-wide CRISPR screen identifies regulators of beta cell function involved in type 2 diabetes risk"

| Gene | Mean log fold change | Effect on insulin content |
|---|---|---|
| AGPAT5 | 2.1342 | Decrease |
| EIF3B | 2.8659 | Decrease |
| RPL12 | 2.052 | Decrease |
| NSF | 2.5825 | Decrease |
| HARS2 | 2.4343 | Decrease |
| NDUFA1 | 1.6306 | Decrease |
| SMC1A | 2.8331 | Decrease |
| WDR35 | 1.9029 | Decrease |
| EP300 | 1.4709 | Decrease |
| MYO9A | 2.6802 | Decrease |
| PARVA | 1.9827 | Decrease |
| GRIP1 | 1.8978 | Decrease |
| HEXIM2 | 1.3545 | Decrease |
| PTP4A1 | 2.6074 | Decrease |
| AIMP2 | 1.0068 | Decrease |
| PSMC4 | 1.9191 | Decrease |
| TP53BP2 | 1.2876 | Decrease |
| HDGF | 1.2184 | Decrease |
| FKBP14 | 1.5067 | Decrease |
| TAP1 | 1.105 | Decrease |
| MARCH3 | 1.7875 | Decrease |
| SH3GL2 | 2.1253 | Decrease |
| KIAA1524 | 1.5713 | Decrease |
| SYDE2 | 1.8711 | Decrease |
| ZNF101 | 1.8076 | Decrease |
| PRKAB1 | 1.6705 | Decrease |
| IFT88 | 1.699 | Decrease |
| YPEL5 | 1.4154 | Decrease |
| ARAP1 | 1.7485 | Decrease |
| C3orf70 | 1.4214 | Decrease |
| FAM184A | 1.3972 | Decrease |
| PPP1R15B | 1.384 | Decrease |
| SLC2A2 | 0.86881 | Decrease |
| AAAS | 2.1018 | Decrease |
| GLCE | 1.8801 | Decrease |
| IKBKG | 1.3903 | Decrease |
| MOB1B | 2.1963 | Decrease |
| VAMP4 | 1.5228 | Decrease |
| SLC7A5 | 1.0087 | Decrease |
| MB21D2 | 1.2686 | Decrease |
| MIPEP | 2.0839 | Decrease |

| | | |
|---|---|---|
| CASC5 | 1.9305 | Decrease |
| SEMA4F | 1.2983 | Decrease |
| CYB561D2 | 0.7521 | Decrease |
| TUBE1 | 1.5298 | Decrease |
| AMER3 | 1.5066 | Decrease |
| RBBP6 | 1.4674 | Decrease |
| DPH1 | 1.8024 | Decrease |
| MAGOH | 2.3212 | Decrease |
| FAM175B | 0.91526 | Decrease |
| DGAT1 | 0.96231 | Decrease |
| NKX2-2 | 1.2148 | Decrease |
| DERL2 | 1.2421 | Decrease |
| FBXL2 | 0.98993 | Decrease |
| RTN3 | 1.4947 | Decrease |
| FAM213A | 1.6758 | Decrease |
| SMTN | 1.5858 | Decrease |
| MBNL1 | 1.7691 | Decrease |
| FERMT2 | 1.6388 | Decrease |
| SLC7A1 | 1.6373 | Decrease |
| PIBF1 | 1.5601 | Decrease |
| TMEM86B | 1.3145 | Decrease |
| NPY | 0.91338 | Decrease |
| PGBD4 | 1.3951 | Decrease |
| ATP5I | -0.82892 | Increase |
| COL1A1 | 1.1399 | Decrease |
| TRIM39 | 1.9988 | Decrease |
| GPSM2 | 1.9117 | Decrease |
| LAMC1 | 1.5196 | Decrease |
| TMEM256 | 1.4527 | Decrease |
| WASF2 | 0.7579 | Decrease |
| BAZ2B | 1.0319 | Decrease |
| PEX13 | 1.7065 | Decrease |
| CNBP | 1.314 | Decrease |
| PRSS23 | 1.4067 | Decrease |
| CALM3 | 1.3306 | Decrease |
| SLC25A19 | 2.0432 | Decrease |
| VIL1 | 1.7482 | Decrease |
| DUSP4 | 1.1443 | Decrease |
| USP45 | 1.2423 | Decrease |
| DHRS1 | 1.5071 | Decrease |
| ASCC3 | 1.4471 | Decrease |
| PPP2R3A | 1.6312 | Decrease |
| POLD2 | 1.3455 | Decrease |

| | | |
|---|---|---|
| SLC25A12 | 1.5179 | Decrease |
| C9orf37 | 1.7719 | Decrease |
| SLC6A8 | 1.6784 | Decrease |
| GLTSCR1L | 1.2624 | Decrease |
| ZBTB22 | 1.129 | Decrease |
| C11orf21 | 1.0255 | Decrease |
| REXO2 | 0.99522 | Decrease |
| VPS37A | 1.5635 | Decrease |
| RDX | 1.155 | Decrease |
| PBK | 1.2888 | Decrease |
| TNPO3 | 1.5661 | Decrease |
| AGPS | 1.345 | Decrease |
| SLMAP | 1.6645 | Decrease |
| TAF7 | 0.84945 | Decrease |
| SRRM1 | 1.1466 | Decrease |
| IRS2 | 1.5712 | Decrease |
| SIPA1L2 | 1.028 | Decrease |
| TCEAL4 | 1.5823 | Decrease |
| AC018755.1 | 1.1706 | Decrease |
| ADAMTS5 | 1.1268 | Decrease |
| PKM | 0.97439 | Decrease |
| DCAF13 | 0.71879 | Decrease |
| CPM | 1.2491 | Decrease |
| HDAC2 | 1.4996 | Decrease |
| PLCL2 | 1.276 | Decrease |
| DDX26B | 0.9321 | Decrease |
| ZNF503 | 1.1861 | Decrease |
| KIAA1586 | 1.2218 | Decrease |
| CHD4 | 1.4781 | Decrease |
| SYNJ1 | 1.0896 | Decrease |
| TEX2 | 1.5849 | Decrease |
| TIAM1 | 1.017 | Decrease |
| NME3 | 1.9053 | Decrease |
| SGPP2 | 1.1227 | Decrease |
| SIN3A | 0.92848 | Decrease |
| TBK1 | 1.5607 | Decrease |
| CALY | 1.321 | Decrease |
| DDX56 | 1.1205 | Decrease |
| SLC2A11 | 1.1633 | Decrease |
| ALG2 | 1.3671 | Decrease |
| ZNF608 | 1.0721 | Decrease |
| ZNF846 | 1.5501 | Decrease |
| PRPF18 | 1.2221 | Decrease |

| | | |
|---|---|---|
| PCNT | 1.1755 | Decrease |
| STARD4 | 1.566 | Decrease |
| WDR83 | 0.83506 | Decrease |
| GRB2 | 1.5086 | Decrease |
| USP14 | 1.3398 | Decrease |
| GMEB1 | 1.1591 | Decrease |
| SLC7A6 | 1.7005 | Decrease |
| GPM6A | 1.2941 | Decrease |
| SMIM5 | 0.99776 | Decrease |
| PAFAH1B1 | 0.93597 | Decrease |
| SGTB | 0.91148 | Decrease |
| SOWAHC | 1.3291 | Decrease |
| DUSP10 | 1.5005 | Decrease |
| ZDHHC4 | 1.4873 | Decrease |
| CACNB3 | 1.1979 | Decrease |
| PPP2R3C | 1.4638 | Decrease |
| EIF3D | 0.69635 | Decrease |
| C3orf14 | 1.7556 | Decrease |
| WDFY1 | 0.88962 | Decrease |
| SLC10A3 | 1.0844 | Decrease |
| PDS5A | 1.2116 | Decrease |
| TMEM2 | 1.6888 | Decrease |
| CALCOCO2 | 1.0745 | Decrease |
| BTBD2 | 1.1143 | Decrease |
| SLC27A5 | 1.1958 | Decrease |
| RAB19 | 0.81418 | Decrease |
| UBE4B | 1.1383 | Decrease |
| TIMM23 | 1.3016 | Decrease |
| DEDD2 | 1.2205 | Decrease |
| NUDT14 | 1.4368 | Decrease |
| C17orf59 | 0.96935 | Decrease |
| IER2 | 1.1046 | Decrease |
| LONRF1 | 1.3479 | Decrease |
| SSTR1 | 0.98459 | Decrease |
| SP1 | 1.3022 | Decrease |
| ANXA13 | 1.2352 | Decrease |
| RSPH9 | 0.99825 | Decrease |
| HDDC3 | 0.94673 | Decrease |
| ABCB6 | 1.1039 | Decrease |
| RNF115 | 1.1493 | Decrease |
| INTS6 | 1.1882 | Decrease |
| HAUS2 | 0.42313 | Decrease |
| UBE2C | 1.0638 | Decrease |

| | | |
|---|---|---|
| TMEM30B | 0.91411 | Decrease |
| MCCC1 | 1.5132 | Decrease |
| NIPBL | 1.1695 | Decrease |
| SAMD4A | 1.2922 | Decrease |
| TENM2 | 1.1071 | Decrease |
| F3 | 1.2835 | Decrease |
| FGFR3 | 0.82426 | Decrease |
| EIF2D | 1.176 | Decrease |
| SOCS7 | 0.26225 | Decrease |
| TM2D2 | 1.0332 | Decrease |
| FBXO16 | 1.4911 | Decrease |
| PARN | 1.4909 | Decrease |
| CAPN7 | 1.175 | Decrease |
| DNAJC1 | 0.81111 | Decrease |
| BMPR1A | 0.9917 | Decrease |
| RPL10A | 1.2158 | Decrease |
| F2R | 1.551 | Decrease |
| SLC44A3 | 0.93304 | Decrease |
| EXOSC3 | 1.4281 | Decrease |
| SERPINA6 | 1.1187 | Decrease |
| PTS | 1.3316 | Decrease |
| MPDU1 | 1.0833 | Decrease |
| FPGS | 1.2025 | Decrease |
| PARP12 | -1.0489 | Increase |
| CDH16 | 1.4427 | Decrease |
| CHST8 | 1.1337 | Decrease |
| CREB3 | 0.99825 | Decrease |
| SMURF2 | 1.2739 | Decrease |
| MBD2 | 0.62786 | Decrease |
| ZNF529 | 1.3575 | Decrease |
| COL4A3BP | 1.3318 | Decrease |
| RAI14 | 1.2967 | Decrease |
| MID1 | 1.04 | Decrease |
| CBR4 | 1.1286 | Decrease |
| STAM | 1.4026 | Decrease |
| CNPY2 | 1.1641 | Decrease |
| FRMD5 | 1.2682 | Decrease |
| RPF1 | 1.4827 | Decrease |
| ALM2-AKAP | 1.2701 | Decrease |
| GABRB3 | 1.0736 | Decrease |
| BTN3A1 | 0.79842 | Decrease |
| MED4 | 0.81514 | Decrease |
| POLR2G | 0.85705 | Decrease |

| | | |
|---|---|---|
| GSDMB | 1.4085 | Decrease |
| AGO4 | 1.2207 | Decrease |
| SCRIB | 0.33653 | Decrease |
| ZNF329 | 1.3335 | Decrease |
| SLC45A4 | 0.91072 | Decrease |
| SMARCC1 | 1.048 | Decrease |
| CCDC66 | 1.1758 | Decrease |
| ARL10 | 1.2483 | Decrease |
| SSR2 | 1.0585 | Decrease |
| LMBR1L | 1.0738 | Decrease |
| SRRM3 | 1.0212 | Decrease |
| GPR119 | 0.98309 | Decrease |
| ACTR3B | 1.0464 | Decrease |
| FBXO22 | 1.0946 | Decrease |
| SRD5A3 | 1.2089 | Decrease |
| AP1S1 | 1.2385 | Decrease |
| OTUB1 | 1.2741 | Decrease |
| CDK5RAP3 | 1.0086 | Decrease |
| ARL6IP6 | 1.2329 | Decrease |
| JAKMIP2 | 1.3106 | Decrease |
| RRAGB | 1.0183 | Decrease |
| RNPS1 | 1.3998 | Decrease |
| CPNE4 | 1.115 | Decrease |
| OSBPL7 | 0.96196 | Decrease |
| SERPINH1 | 1.3303 | Decrease |
| EPB41L1 | 0.9267 | Decrease |
| PSEN1 | 1.3355 | Decrease |
| ATP8B2 | 1.1916 | Decrease |
| PELO | 0.87529 | Decrease |
| C12orf29 | 1.1138 | Decrease |
| MVK | 1.453 | Decrease |
| NLK | 1.0001 | Decrease |
| SLC16A10 | 1.1569 | Decrease |
| RBM43 | 1.1936 | Decrease |
| NUP54 | 0.99691 | Decrease |
| PCED1A | 1.0395 | Decrease |
| ZNF510 | 1.2974 | Decrease |
| INPP5J | 0.63313 | Decrease |
| INS | 1.0593 | Decrease |
| GALE | 0.95466 | Decrease |
| CHODL | 1.045 | Decrease |
| TMEM39B | 1.3191 | Decrease |
| TRPM2 | 1.026 | Decrease |

| | | |
|---|---|---|
| CXorf56 | 1.6605 | Decrease |
| DUSP3 | 1.0871 | Decrease |
| KIAA0247 | 1.1512 | Decrease |
| PEX10 | 1.1507 | Decrease |
| FIBP | 1.0112 | Decrease |
| CEP85L | 1.4025 | Decrease |
| ZGLP1 | 0.98705 | Decrease |
| ASXL3 | 1.0081 | Decrease |
| ZNF550 | 0.77412 | Decrease |
| ZBTB24 | 0.89339 | Decrease |
| FFAR1 | 1.2275 | Decrease |
| ARNT2 | 0.5061 | Decrease |
| DLL3 | 1.2887 | Decrease |
| MED11 | 1.0411 | Decrease |
| ANP32E | 1.1211 | Decrease |
| ANXA10 | 0.57976 | Decrease |
| ACO2 | 1.0176 | Decrease |
| ZCCHC4 | 1.1074 | Decrease |
| PSMB2 | 1.1897 | Decrease |
| SLC25A40 | 1.1113 | Decrease |
| ASAP1 | 1.0409 | Decrease |
| FLII | 1.0823 | Decrease |
| ZNF432 | 0.39443 | Decrease |
| LURAP1L | 0.87078 | Decrease |
| SERTAD2 | 1.0889 | Decrease |
| CHTOP | 0.73978 | Decrease |
| MGA | 1.1732 | Decrease |
| ANKRD7 | 0.84834 | Decrease |
| CDK5R2 | 1.1034 | Decrease |
| CLK4 | 0.74779 | Decrease |
| ITM2C | 0.73017 | Decrease |
| CFLAR | 1.1038 | Decrease |
| TRIM45 | 0.66403 | Decrease |
| DHX8 | 1.0213 | Decrease |
| GPS1 | 0.98874 | Decrease |
| TBC1D19 | 0.29839 | Decrease |
| RAP1A | 1.0736 | Decrease |
| DOPEY2 | 0.82147 | Decrease |
| NUP107 | 0.94979 | Decrease |
| JARID2 | 0.33975 | Decrease |
| SMYD5 | 1.2049 | Decrease |
| MMP16 | 1.2611 | Decrease |
| CETN3 | 0.57769 | Decrease |

| | | |
|---|---|---|
| NUDT3 | 1.0947 | Decrease |
| C7orf41 | -0.73144 | Increase |
| UBE2D1 | 1.1993 | Decrease |
| MRPL16 | 1.0243 | Decrease |
| TSTD1 | 0.87149 | Decrease |
| GJD2 | 0.82885 | Decrease |
| EIF3J | 0.77937 | Decrease |
| KALRN | 1.1739 | Decrease |
| QSOX2 | 0.9479 | Decrease |
| WDR7 | 0.75451 | Decrease |
| IREB2 | 0.76342 | Decrease |
| ZNF214 | 1.1781 | Decrease |
| HOMER2 | 0.84087 | Decrease |
| POLM | 1.1791 | Decrease |
| LRRN3 | 1.1311 | Decrease |
| NECAB2 | 0.70199 | Decrease |
| HUWE1 | 0.76028 | Decrease |
| PLK1 | 1.2327 | Decrease |
| MPST | -1.3021 | Increase |
| UBE2J1 | 1.0673 | Decrease |
| PPAP2B | 0.73861 | Decrease |
| C20orf112 | 1.2357 | Decrease |
| DOK1 | 0.79882 | Decrease |
| DPYSL5 | 0.60515 | Decrease |
| SDCCAG8 | 0.70192 | Decrease |
| PALLD | 0.81382 | Decrease |
| RPS6KC1 | 0.99293 | Decrease |
| FTSJ1 | 0.8383 | Decrease |
| RPL18A | 1.2983 | Decrease |
| DDAH2 | 0.76168 | Decrease |
| CDYL2 | 1.0024 | Decrease |
| FBXO48 | 1.0846 | Decrease |
| RBMS1 | 0.9105 | Decrease |
| RAD23A | 0.89095 | Decrease |
| GINS3 | 0.74477 | Decrease |
| WIPF1 | 0.84887 | Decrease |
| KANK1 | 0.96197 | Decrease |
| GALNT7 | 1.1398 | Decrease |
| MED13L | 0.86581 | Decrease |
| ZNF821 | 1.0192 | Decrease |
| PITPNM2 | 0.99058 | Decrease |
| IFNGR2 | 0.83005 | Decrease |
| NDUFAF2 | 1.0367 | Decrease |

| | | |
|---|---|---|
| NUPL1 | 0.96487 | Decrease |
| NDUFAF4 | 0.84558 | Decrease |
| MESDC2 | 1.1164 | Decrease |
| EXOSC9 | 1.2793 | Decrease |
| NCOA6 | 1.0964 | Decrease |
| NCOA4 | 0.97316 | Decrease |
| MAFG | 0.98148 | Decrease |
| TNFRSF11A | 1.0158 | Decrease |
| AGPAT2 | 0.53893 | Decrease |
| TCF19 | 0.69866 | Decrease |
| IGF2 | 0.85199 | Decrease |
| TAGLN | 0.38694 | Decrease |
| ELAVL1 | 0.84406 | Decrease |
| UBR1 | 1.1283 | Decrease |
| C12orf52 | 0.52356 | Decrease |
| MAPK1 | 1.0446 | Decrease |
| DDX23 | 0.79136 | Decrease |
| ZNF524 | 1.0125 | Decrease |
| FGFR1OP | 1.0716 | Decrease |
| RBP1 | 1.0669 | Decrease |
| COA5 | 0.64784 | Decrease |
| C7orf60 | 1.0325 | Decrease |
| BTBD10 | 1.0319 | Decrease |
| TMEM132D | 0.85866 | Decrease |
| ZNF654 | 0.83762 | Decrease |
| BTF3L4 | 0.86571 | Decrease |
| ZCCHC9 | 0.98358 | Decrease |
| ADNP | 0.9912 | Decrease |
| PRKD1 | 0.875 | Decrease |
| HSPA8 | 0.81331 | Decrease |
| TMEM68 | 0.90746 | Decrease |
| PCGF5 | 0.85942 | Decrease |
| SAE1 | 0.94922 | Decrease |
| MRPS25 | 1.0502 | Decrease |
| DHX36 | 0.82639 | Decrease |
| ATP6V1C1 | 0.66221 | Decrease |
| PDE4D | 1.1062 | Decrease |
| LAS1L | 0.96268 | Decrease |
| MRPL28 | 0.94149 | Decrease |
| HMGXB4 | 0.88796 | Decrease |
| HNRNPDL | 1.0125 | Decrease |
| SLC29A4 | 0.45219 | Decrease |
| DCUN1D4 | 1.3779 | Decrease |

| | | |
|---|---|---|
| DPCD | 1.0447 | Decrease |
| KREMEN1 | 0.39301 | Decrease |
| METTL5 | 0.5093 | Decrease |
| SLC35G2 | 0.90109 | Decrease |
| MED18 | 0.84647 | Decrease |
| PTPN1 | 1.1306 | Decrease |
| ZCCHC2 | 0.50546 | Decrease |
| LOXL4 | 0.52638 | Decrease |
| CRB3 | 0.77113 | Decrease |
| PSME4 | 0.83049 | Decrease |
| PHACTR2 | 0.76559 | Decrease |
| DPYSL2 | 0.95314 | Decrease |
| RCC2 | 0.98031 | Decrease |
| GNL3 | 0.78882 | Decrease |
| ABHD8 | 0.38544 | Decrease |
| CMTM6 | -0.23869 | Increase |
| ANAPC5 | 0.80375 | Decrease |
| RPL10 | 1.0426 | Decrease |
| FAM212B | 0.94091 | Decrease |
| TBC1D4 | 0.77573 | Decrease |
| ANO6 | 1.0706 | Decrease |
| NUP188 | 0.78533 | Decrease |
| DOK4 | 0.75297 | Decrease |
| ZNF141 | 0.71702 | Decrease |
| CIAO1 | 0.4643 | Decrease |
| ARF1 | 0.55193 | Decrease |
| ZNF155 | 0.71369 | Decrease |
| FAM109A | 0.80655 | Decrease |
| C14orf39 | 0.98803 | Decrease |
| TOMM6 | 0.47169 | Decrease |
| ZNF24 | 0.65659 | Decrease |
| PPP6R3 | 0.79534 | Decrease |
| TAF3 | 0.66582 | Decrease |
| HIST1H2BK | 0.99609 | Decrease |
| TAF8 | 0.91843 | Decrease |
| RPUSD2 | 1.0568 | Decrease |
| MRPS24 | 0.85085 | Decrease |
| FBXO21 | 0.8411 | Decrease |
| PRKAA1 | 0.92974 | Decrease |
| IPO4 | 0.67357 | Decrease |
| CAT | 1.2751 | Decrease |
| FBXW8 | -0.33827 | Increase |
| IQSEC1 | 0.5309 | Decrease |

| | | |
|---|---|---|
| QDPR | 0.62851 | Decrease |
| USP6NL | 0.7306 | Decrease |
| DNAJB7 | 0.45177 | Decrease |
| 9-Mar | 0.80004 | Decrease |
| ARMCX5 | 0.83118 | Decrease |
| TMX4 | -0.75754 | Increase |
| TSPAN2 | 0.86282 | Decrease |
| PRDM10 | 0.70835 | Decrease |
| HM13 | 0.67128 | Decrease |
| TSHZ2 | 0.86652 | Decrease |
| ARL14EP | 0.64244 | Decrease |
| COX19 | 0.87764 | Decrease |
| FDX1 | 0.91154 | Decrease |
| ASH2L | 0.6178 | Decrease |
| ANTXR2 | 0.32543 | Decrease |
| MTIF2 | 0.85522 | Decrease |
| RBM19 | 0.88679 | Decrease |
| PPAPDC2 | 0.73519 | Decrease |
| NOL8 | 0.6942 | Decrease |
| RPUSD3 | 0.83847 | Decrease |
| PPA2 | 0.65904 | Decrease |
| CDK5RAP2 | 0.96184 | Decrease |
| CTNNA2 | 0.84745 | Decrease |
| ZFP30 | 1.0008 | Decrease |
| SFR1 | 0.55777 | Decrease |
| SLC8B1 | 0.71096 | Decrease |
| ZNF684 | 0.97435 | Decrease |
| CHRNA5 | 0.776 | Decrease |
| WDR27 | 0.39713 | Decrease |
| C2orf76 | 0.6225 | Decrease |
| TRIM41 | 0.86348 | Decrease |
| AURKA | 0.69006 | Decrease |
| GPR137 | 0.47045 | Decrease |
| MLH3 | 0.9159 | Decrease |
| FLNB | 0.42319 | Decrease |
| C7orf49 | 0.57614 | Decrease |
| SON | 1.0395 | Decrease |
| TIGD2 | 0.76883 | Decrease |
| MTERFD1 | 0.74006 | Decrease |
| DUSP18 | 0.36615 | Decrease |
| ZNF426 | 0.39117 | Decrease |
| DNAJC4 | 0.64778 | Decrease |
| SHQ1 | 0.70085 | Decrease |

| | | |
|---|---|---|
| MED23 | 0.58596 | Decrease |
| NAA40 | 0.77757 | Decrease |
| TMEM144 | 0.65704 | Decrease |
| BLCAP | 0.46821 | Decrease |
| SPATA24 | 0.13191 | Decrease |
| OSBP2 | 0.20825 | Decrease |
| ANKRD46 | -0.87669 | Increase |
| AP5S1 | 0.17943 | Decrease |
| GEMIN6 | 0.52253 | Decrease |
| FAM172A | 0.60914 | Decrease |
| NPHP4 | 0.46499 | Decrease |
| TAPBPL | 0.25611 | Decrease |
| DVL2 | 0.36616 | Decrease |
| IL1R2 | 0.49474 | Decrease |
| GSN | 0.70791 | Decrease |
| PRPF38A | 0.31695 | Decrease |
| COL4A1 | 0.33467 | Decrease |
| TMEM251 | 0.56511 | Decrease |
| USP12 | 0.30632 | Decrease |
| ZNF616 | 0.64281 | Decrease |
| RUFY1 | 0.66453 | Decrease |
| DDX3Y | 0.46491 | Decrease |
| FBXO46 | 0.60517 | Decrease |
| ZNF404 | 0.33025 | Decrease |
| TTYH3 | 0.34258 | Decrease |
| RFWD2 | 0.8621 | Decrease |
| DMTN | 0.64018 | Decrease |
| RSPRY1 | 0.77445 | Decrease |
| SMC3 | 0.4149 | Decrease |
| GPR98 | 0.15947 | Decrease |
| FAM178A | 0.67494 | Decrease |
| HBS1L | 0.6491 | Decrease |
| IKZF4 | 0.5831 | Decrease |
| TPP1 | 0.54622 | Decrease |
| NRAS | 0.34705 | Decrease |
| RSRC1 | 0.50227 | Decrease |
| GEMIN8 | 0.40602 | Decrease |
| GLI4 | 0.5022 | Decrease |
| GBP3 | -0.60088 | Increase |
| WDR81 | 0.51694 | Decrease |
| RND3 | 0.72533 | Decrease |
| PCDHA2 | 0.48315 | Decrease |
| ARIH2 | 0.29702 | Decrease |

| | | |
|---|---|---|
| PLA2G4C | 0.42569 | Decrease |
| RAD52 | 0.72593 | Decrease |
| SLC7A2 | 0.4927 | Decrease |
| FUT4 | 0.26972 | Decrease |
| ZNF259 | 0.52581 | Decrease |
| IBTK | 0.48759 | Decrease |
| NISCH | 0.19543 | Decrease |
| MRPL19 | 0.33484 | Decrease |
| LCAT | -1.606 | Increase |
| DOCK7 | 0.50189 | Decrease |
| FSTL3 | 0.46989 | Decrease |
| TEX10 | 0.55276 | Decrease |
| C1orf198 | -1.3921 | Increase |
| SLC25A43 | 0.20626 | Decrease |
| PANK3 | 0.39243 | Decrease |
| SLC25A39 | -0.48829 | Increase |
| FADS1 | -1.5294 | Increase |
| PGLS | -2.3068 | Increase |
| GPAA1 | -1.5067 | Increase |
| HNRNPM | -1.8841 | Increase |
| PLD3 | -1.5006 | Increase |
| ACTN1 | -1.5379 | Increase |
| RAB11FIP3 | -1.6839 | Increase |
| RELB | -1.455 | Increase |
| SCO2 | -1.7642 | Increase |
| CARHSP1 | -1.3506 | Increase |
| SIDT2 | -1.5989 | Increase |
| SQSTM1 | -1.6914 | Increase |
| C21orf2 | -1.293 | Increase |
| TLDC1 | -1.4157 | Increase |
| MRPL53 | -1.1815 | Increase |
| NAV1 | -1.1381 | Increase |
| TNFAIP8L1 | -1.3476 | Increase |
| RPUSD1 | -1.6244 | Increase |
| PHF8 | -1.6143 | Increase |
| SMURF1 | -1.4531 | Increase |
| SLC25A10 | -1.3512 | Increase |
| LRRC61 | -1.2335 | Increase |
| POP4 | -1.4278 | Increase |
| PSAP | -1.3824 | Increase |
| AGO3 | -0.91766 | Increase |
| NOC2L | -1.0883 | Increase |
| PCMT1 | -1.2769 | Increase |

| | | |
|---|---|---|
| XYLT2 | -1.0793 | Increase |
| AGO1 | -0.82049 | Increase |
| B4GALNT4 | -1.2548 | Increase |
| LETMD1 | -1.3124 | Increase |
| ANKRD13D | -0.89667 | Increase |
| ZNF672 | -1.2563 | Increase |
| PDCD11 | -0.99851 | Increase |
| SLX4 | -1.3399 | Increase |
| GATAD1 | -0.86942 | Increase |
| MIF | -0.69335 | Increase |
| RPL35 | -0.91465 | Increase |
| LHFP | -0.86672 | Increase |
| PBX3 | -1.3498 | Increase |
| ADCY6 | -1.4016 | Increase |
| GJD3 | -0.77409 | Increase |
| FAM46C | -1.0806 | Increase |
| ORAOV1 | -0.79123 | Increase |
| PRKCD | -1.017 | Increase |
| POLH | -0.64276 | Increase |
| RAB14 | -0.59583 | Increase |
| CSTF2 | -0.087204 | Increase |
| ATP5H | -1.1225 | Increase |
| EID2B | -0.82751 | Increase |
